# A Deep-Learning Atlas of XPO1-Mediated Nuclear Export at Proteome Scale

**DOI:** 10.64898/2026.03.25.713363

**Authors:** Sajina Dhungel, Sihara de Zoysa, DeAnna Burns, Linden McGregor, RP Rajesh, Ridila Alam, Diya Arain, Vivaan Bhaskar, Jayden Jeong, Aarush Kikani, Eashan Kolli, Zain Mardini, Aanya Parasramka, Eden Potterton, Saachi Thomas, Chintan K. Kikani

**Affiliations:** University of Kentucky, Department of Biology, Lexington, KY, USA; University of South Carolina; Paul Laurence Dunbar High School, Lexington, KY, USA

**Author notes:** Corresponding author: Chintan K. Kikani, Department of Biology, University of Kentucky, Lexington, KY, USA.

## Abstract

Exportin 1 (XPO1/CRM1) is the principal nuclear export receptor for cargos bearing hydrophobic nuclear export sequences (NESs). Dysregulation of XPO1-dependent export is implicated in cancer, neurodegeneration, and other diseases, yet a comprehensive view of XPO1 function remains limited by the poor reliability of sequence-based NES prediction. Existing predictors are largely derived from a small set of XPO1-cargo structures and are therefore biased toward canonical docking geometries, limiting their ability to detect NESs that engage XPO1 through noncanonical pocket-occupancy patterns. We hypothesized that deep learning-based structural modeling could overcome this limitation by directly sampling binding geometries. Using AlphaFold 3, we modeled full-length cargo-XPO1-RanGTP complexes for more than 4,000 human proteins and identified over 3,000 previously uncharacterized, high-confidence NESs. Integration of AlphaFold predictions with unsupervised structural geometry analysis and experimental validation identified both canonical NESs and noncanonical sequence patterns exhibiting atypical anchor-residue usage, expanding the structural language of XPO1-recognized NESs. Groove-resolved contact maps further revealed helix rotation within the export groove as a regulatory feature that can rewire pocket usage without altering the core NES sequence, enabling PTM- and cofactor-sensitive tuning of export strength. This exportome atlas resolves many previously ambiguous or unidentified NESs in disease-associated proteins and across major cellular systems, including centrosome organization, mRNA processing, ubiquitin signaling, kinase networks, ribosome quality control, and macroautophagy. We further identified recurrent NES-NLS tandem motifs encoded in primary sequence, suggesting coordinated regulation of nucleocytoplasmic transport. Together, our deep learning–based exportome atlas, integrated with NLS maps and accessible through a web-searchable resource, defines an expanded and regulatable code of nuclear transport at proteome scale and offers a framework for dissecting nuclear trafficking and its dysregulation in human disease.

## INTRODUCTION

Macromolecular traffic between the nucleus and cytoplasm is carried out by karyopherin transport receptors that read short peptide or structural motifs on cargo^1^. Importin and exportin proteins shuttle proteins across the nuclear pore via the Ran-dependent guanosine triphosphate (GTP)-guanosine diphosphate (GDP) cycle^2^. The exportin family includes Exportin 1 (XPO1), CSE1L/XPO2, XPO4, XPO5, XPO6, XPO7, and XPOT (exportin-t), which shuttle distinct classes of cargos out of the nucleus. For example, XPO2 recycles importin-α, XPOT exports mature tRNAs, XPO5 exports pre-miRNA hairpins and other structured RNAs, XPO6 exports profilin-actin complexes, and XPO4 exports eIF5A and Smad3, and in specific contexts, mediates the import of Sox2/SRY. XPO7 exports a broad range of proteins with context-dependent selectivity. In contrast, XPO1 primarily engages with leucine-rich Nuclear Export Sequence (NES)-bearing proteins and is therefore the dominant mediator of protein nuclear export in mammalian cells^3–9^.

XPO1 is an evolutionarily conserved gatekeeper of cellular homeostasis in multicellular organisms, as it controls the nuclear-cytoplasmic partitioning of proteins involved in the cell cycle, DNA damage response, and key transcriptional and translational effectors in response to signaling cues^10,11^. XPO1 is frequently overexpressed in malignancies, and higher expression correlates with advanced stage and poor survival, consistent with the dependence of cancer cells on high nuclear export flux^12,13^. Pharmacological inhibition with the selective nuclear export inhibitor Selinexor has clinically validated this dependency in multiple myeloma and relapsed or refractory diffuse large B-cell lymphoma (DLBCL)^14,15^.

As interactors of XPO1, NESs are often described as short leucine-rich linear motifs; however, their recognition by XPO1 is intrinsically structural^16^. A productive XPO1-NES interaction requires a series of hydrophobic anchor residues that engage XPO1 while maintaining polar contacts that ensure specificity. Structurally, these anchors dock into a conserved NES-binding groove on the inner surface of the XPO1 toroid, formed by 20 HEAT repeats (H1-H20), each comprising paired antiparallel α-helices. NES engagement centers on HEAT repeats 11 and 12 (H11 and H12, Figure S1A), where a series of hydrophobic pockets, P0-P4, accommodate four to five hydrophobic anchor positions, Φ0-Φ4, from the cargo^17,18^. Despite the diversity of NES backbone geometries, ranging from extended strands to multi-turn helices, the spatial engagement of these anchors was preserved. Binding can occur in the canonical plus orientation, N-to-C, Φ0→Φ4, or in a reverse, minus orientation, C-to-N, Φ4→Φ0, effectively doubling the allowable solutions and expanding the classification to at least eleven consensus classes^19^. A conserved lysine on human XPO1, Lys568, contacts NES backbone atoms and acts as a geometry and sequence screener that helps discriminate functional NESs from NES-like stretches that fail to achieve the correct three-dimensional presentation^16,19^. Thus, NES engagement with XPO1 is exquisitely encoded by the structural arrangement of the cargo peptide.

Conventional low-throughput NES mapping combines biochemical analysis, nuclear exportability of peptide-linked fluorescence reporters, and point mutational analysis of full-length proteins in a biological context to identify functional NESs. Despite extensive efforts to define the XPO1 cargo repertoire, many nuclear export signals (NESs) remain undetected because of structural and contextual constraints. Sequence-based identification relying on hydrophobic spacing motifs often overpredicts NESs, as many candidates fail to adopt the amphipathic α-helical conformation or side-chain orientation necessary to engage the XPO1 P0-P4 binding pockets. Additionally, a substantial subset of NESs is conditionally formed, requiring disorder-to-order transition upon XPO1 engagement. Additionally, functional discrepancies are frequently observed between fluorescence-based peptide reporter assays and full-length protein architecture because, in native protein contexts, these motifs may be sterically occluded by intramolecular packing or masked through interactions with binding partners. These factors collectively contribute to the limitations of sequence-only scans and interactome assays, which tend to overcall false positives and miss genuine context-dependent cargos.

Thus, accurate identification of NESs remains a longstanding challenge in biology. The interaction between XPO1 and NESs highlights both the limitations of traditional sequence-based prediction methods and the promise of advanced structure-based modeling to identify previously unseen motif-receptor interactions with high fidelity. In this study, we harness recent advances in deep-learning-based modeling driven by AlphaFold 3.0 to construct complete export complexes comprising XPO1, Ran-GTP, and cargo across more than 4,000 human proteins. We developed a structure-based NES detection workflow that scores and classifies candidate sequences based on their bound-state structural geometry. Our comprehensive analysis and validation not only recovered previously characterized NESs but also identified over 3,000 high-confidence candidates, collectively expanding the NES exportome across diverse functional and enzymatic protein classes. Intriguingly, many of these newly elucidated NESs were found adjacent to nuclear localization signals (NLSs), suggesting a potential mechanism for switch-like regulation within the nucleocytoplasmic transport system.

## Results

### AF3 consistently models experimentally validated NES motifs across diverse cargo proteins

To benchmark AF3’s capacity to accurately recover experimentally determined NES structures, we modeled well-characterized NES-containing proteins in their full-length contexts. AF3 predictions were generated in template-free mode as XPO1-RanGTP-RanBP1-cargo complexes, with sequences matching those of the Exportin-1:NES cargo structure (PDB: 6CIT)^19^. AF3 consistently produced high-fidelity structural models for representative NES classes (Figure S1B) and accurately captured the canonical engagement of NESs within the XPO1 binding groove between helices H11 and H12 for both forward (Figure S1C) and reverse (Figure S1D) oriented NES. Notably, backbone RMSDs for modeled NES peptides were consistently below 1.0 Å relative to their corresponding experimental structures, underscoring AF3’s ability to resolve motif-receptor interactions with high accuracy.

Next, we investigated a set of proteins with experimentally characterized or proposed NES motifs that lack resolved structural data. These cargo proteins span diverse functional categories, including kinases, cell cycle regulators, tumor suppressors, and phosphatases, providing a rigorous test of AF3’s ability to accurately identify and position validated NESs within the conserved exportin groove formed by HEAT repeats 11-12 of XPO1.

Among these, we first modeled functional NES in 3′-Phosphoinositide-Dependent Kinase 1 (PDPK1, also known as PDK1). Mutagenesis experiments demonstrated that residues 373-385, located between the kinase and PH domains of mouse Pdpk1, are essential for its nuclear export (Figure 1A). Specifically, substitution of two key hydrophobic residues, Leu^383^ and Phe^386^ (Leu^380^ and Phe^383^ for human PDPK1), was sufficient to block export and retain PDPK1 in the nucleus, confirming their critical role in NES function^20^. Consistent with these findings, AF3 reliably modeled residues 376-388 of human PDPK1 (^376^YDNLLSQFGCMQV^388^) as an α-helix positioned within the XPO1 binding groove (Figure 1B). The predicted complex revealed that L^380^, F^383^, M^386^, and V^388^ are embedded in the P1-P4 hydrophobic pockets of the XPO1 H11-H12 groove, respectively (Figure 1C). Additional stabilization was provided by K^579^ (equivalent to human XPO1-K568, Figure S1E), which formed hydrogen bonds with backbone carbonyls along the helix, and by acidic residues E^540^ and E^586^ of XPO1, which interacted with the NES peptide as it exited the groove (Figure 1C). Collectively, these features recapitulate the canonical Clas1a NES occupation within XPO1 groove and underscore AF3’s capacity to capture previously unseen NES-XPO1 interactions with high accuracy.

**Figure 1.**
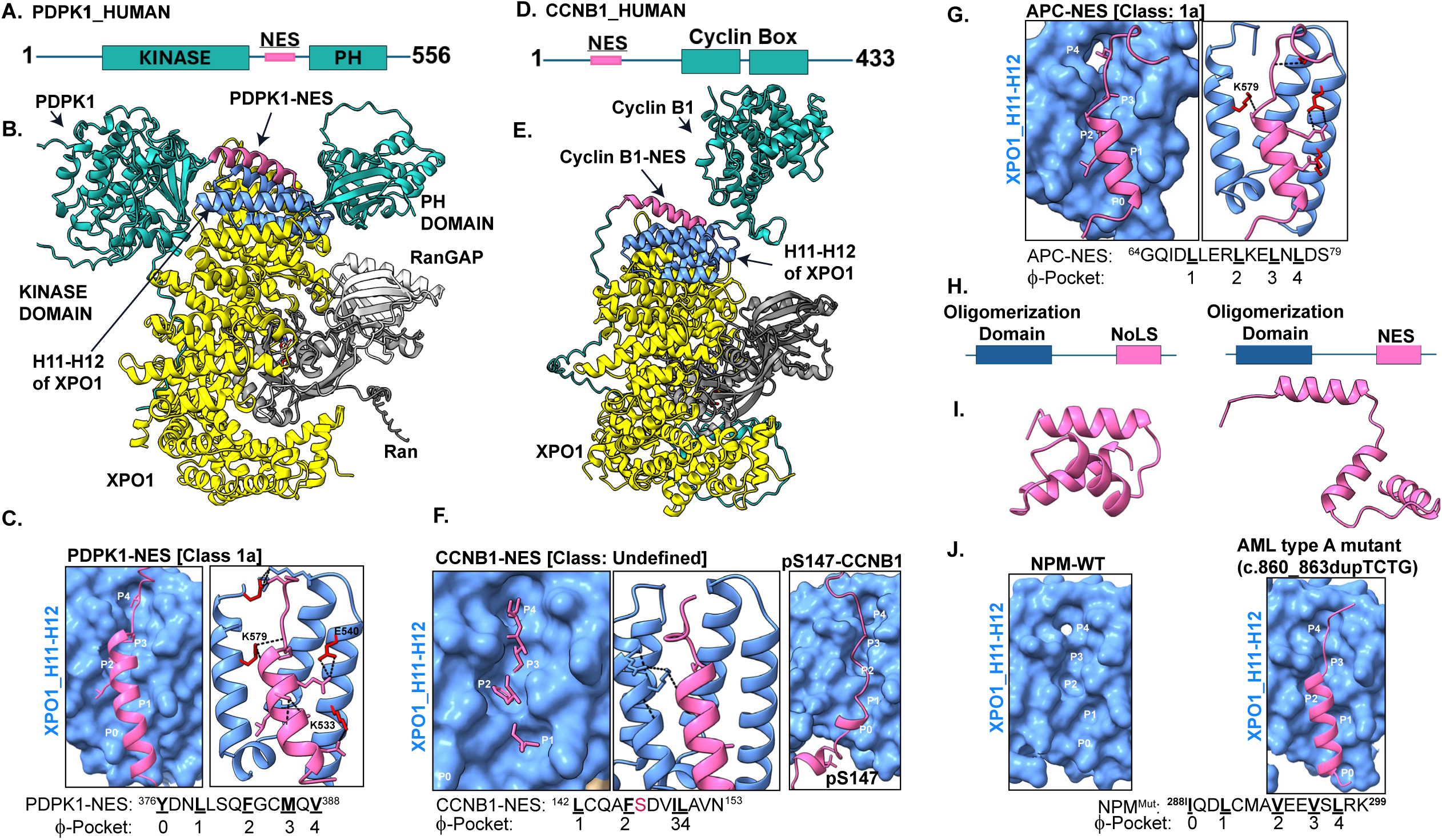
Canonical pocket-resolved binding of unseen leucine-rich NESs in the XPO1 H11-H12 groove by AlphaFold 3.0. **(A)** Schematic domain organization of PDPK1 showing the experimentally validated NES between the N-terminal kinase and C-terminal PH domains. **(B)** Overall AF3 model of the human export complex with XPO1 (yellow), Ran-GTP (dark gray), RanBP1 (light gray), and PDPK1 (teal). The PDPK1 NES helix (magenta) docks in the H11-H12 groove (blue). **(C)** PDPK1-NES bound to XPO1: left, surface rendering of the groove with hydrophobic pockets P0-P4 labeled; right, ribbon view highlighting polar contacts to the selectivity-filter lysine (yeast numbering K579; equivalent to human K568) and neighboring residues. Bottom, PDPK1 NES sequence (^376^YDNLLLSQFCGMQV^388^) with Φ-positions mapped to pockets 0-4. **(D)** Schematic domain organization of Cyclin B1 (CCNB1) with the NES N-terminal to the Cyclin box. **(E)** AF3 model of XPO1-Ran•GTP-RanBP1 bound to CCNB1 (teal) showing NES engagement in H11-H12 (magenta helix). **(F)** CCNB1-NES bound to XPO1: left, surface view of pocket occupancy P1-P4 by NES residues in non-canonical spacing pattern (3-3-0); middle, ribbon views indicating the geometric relationship of the NES helix to K579; right, Phosphorylation of CCNB1 at pS147 disrupts NES-Helix engagement and orientation with XPO1. Bottom, CCNB1 NES sequence (^142^LCQAFESDVILAVN^153^) annotated to occupy non-canonical pocket register 1-4. Ser147, the target of mitotic PTM, is highlighted in red. **(G)** APC NES (^64^GQIDLLERLKELNLDS^79^) in the XPO1 groove: surface (left) and ribbon (right) views depicting Φ-side-chain placement across P0-P4 and the proximity to K579. **(H)** NPM1 constructs: wild-type with N-terminal oligomerization domain and C-terminal NoLS (left) and AML type-A mutant (c.860_863dupTCTG) generating a C-terminal NES (right). **(I)** Predicted conformations of the WT (left) and AML-mutant (right) C-terminal segments illustrating helix formation in the mutant. **(J)** XPO1 surface views for NPM1 peptides: WT segment does not productively occupy P0-P4, whereas the AML mutant peptide binds in the canonical register; sequence shown with pocket assignment.

Cyclin B1 contains a PTM-regulated leucine-rich nuclear export sequence (140-153) embedded within its N-terminal cytoplasmic retention sequence (CRS), which drives CRM1/XPO1-dependent export and underlies the predominantly cytoplasmic localization of Cdk1-Cyclin B1 during S and G2(Figure 1D)^21^. In our AF3 models, residues 142-151 of Cyclin B1 are consistently positioned within the H11-H12 NES-binding groove of XPO1; however, the helix engages the groove using a non-canonical set of hydrophobic anchors in ^142^**<u>L</u>**CQA**<u>F</u>**SDV**<u>IL</u>**AV^153^, arranged in a 3-3-0 spacing pattern, rather than the predicted 3-2-1 spacing of a canonical Class 1a NES, featuring anchors ^142^**<u>L</u>**CQA**<u>F</u>**SD**<u>V</u>**<u>I**L**</u>AV^153^ (Figure 1D-E). This alternative anchor usage is reproducible across AF3 runs, appearing in 4 of 5 models and remaining unchanged with different modeling seeds. Because of its non-canonical spacing, we classified the Cyclin B1 NES as undefined (Figure 1F) (see discussion).

Cyclin B1 is phosphorylated within its CRS at the onset of mitosis by PLK1 at Ser147, a modification that disrupts XPO1 binding and leads to nuclear retention^22^. AF3 modeling of this PTM illustrates how phosphorylation can modulate export by reshaping helix presentation in the groove. When Ser147 is modeled as phosphoserine, the NES helix kinks and partially unravels around the modified residue, and the NES segment swings away from the groove surface. In these phospho-NES models, the phosphoserine side chain protrudes toward XPO1, the Φ-pocket register is lost, and the NES helix is effectively mis-localized out of the groove, providing a structural explanation consistent with the observed phosphorylation-dependent loss of Cyclin B1 export (Figure 1F, right)^21^.

In the tumor suppressor APC, which regulates β-catenin degradation, multiple NESs have been reported at both the N- and C-termini; however, their relative contributions to nuclear export remain unclear. Early studies identified two N-terminal NESs (residues 68-77 and 165-174), both capable of driving export in reporter assays, while additional C-terminal motifs have been proposed^23^. Yet, structural and biochemical evidence have cast doubt on several of these candidates, as key hydrophobic residues appear buried in available structures or fail to bind XPO1 fragments in vitro.

AF3 consistently modeled only NES1 (residues ^67^LLERLKELNL^78^) as an α-helix inserted into the canonical XPO1 groove formed by HEAT repeats 11-12 (Figure 1G). In this model, hydrophobic residues L68, L69, L72, and L75 occupy pockets P1-P3, while acidic side chains form polar contacts with basic residues of XPO1, including K567 and K579 (Figure 1G). In contrast, other reported motifs did not engage the groove in AF3 predictions, suggesting that NES1 likely functions as a constitutive export signal. Additional sites may be conformationally masked or regulated through partner interactions, highlighting the complexity of NES accessibility and function in APC.

To evaluate AF3’s ability to model disease-associated mutations that alter nucleocytoplasmic transport, we examined nucleophosmin (NPM), a protein frequently mislocalized in acute myeloid leukemia (AML) due to recurrent frameshift mutations^24^. In the wild-type protein, a C-terminal nucleolar localization sequence (NoLS) enforces nucleolar retention and includes a weak nuclear export signal (NES) at its N-terminus (Figure 1H). Reflecting this limited export capacity, AF3 did not consistently model NES-compatible helices that could engage XPO1 in wild-type NPM1 structures. In contrast, AML type A frameshift mutations (e.g., c.860_863dupTCTG) disrupt the NoLS and simultaneously generate a leucine-rich C-terminal tail (Figure 1H). The predicted complex revealed a conformational rearrangement within the mutated NoLS domain that exposes this tail as an NES-compatible helix, positioning it for productive engagement with XPO1 (Figure 1I). AF3 consistently modeled this mutant segment as an amphipathic α-helix inserted into the NES-binding groove of XPO1, with hydrophobic residues aligned across pockets P0-P4 and stabilizing polar contacts along the peptide backbone (Figure 1J). This clear structural distinction between wild-type and mutant forms highlights AF3’s capacity to capture gain-of-function NESs arising from pathogenic mutations. Broadly, it demonstrates the utility of deep learning-based structural modeling for mechanistically linking sequence variants to altered nucleocytoplasmic trafficking and functional outcomes.

Taken together, these examples, spanning constitutive, PTM-regulated, and disease-linked NES detection, demonstrate that AF3 can consistently identify and correctly position NES motifs across a wide range of cargo proteins. The models reproduce the hallmark features of NES recognition, even in the full-length protein context, including the stepwise insertion of hydrophobic side chains into the P0-P4 pockets, salt-bridge formation at the groove entrance, and helix stabilization by K579 and other basic residues of XPO1. Beyond identifying canonical NES engagement, full-length models provide a framework for understanding and visualizing how NES presentation is governed by the structural context of the cargo protein, revealing the experimentally testable conformational reorganization required to expose the motif and enable productive engagement with XPO1.

### Proteome-wide NES discovery workflow successfully identifies hundreds of novel nuclear export sequences across functional protein families

To extend AF3 modeling of XPO1-cargo interactions toward proteome-wide NES discovery, we assembled a dataset of human proteins implicated in XPO1-dependent nuclear export but lacking experimentally defined NES. We compiled an initial set of ∼1,500 high-confidence XPO1 interactors from curated BioGRID datasets^25,26^ (Table S1). This list was expanded by incorporating candidate nucleocytoplasmic shuttling proteins identified through subcellular localization data from the Human Protein Atlas (HPA)^27^ and UniProt, retaining cargos with dual nuclear and cytoplasmic distribution, including those localized in cytoplasmic organelles. We further prioritized proteins of biomedical relevance, including those represented in Pan-Cancer biomarker diagnostic panels^28^, enriched in disease-associated gene sets, or belonging to critical signaling classes such as E3 ligases, serine/threonine kinases, and metabolic enzymes. This integrative curation yielded a final set of ∼4000 proteins for structural modeling and NES identification in the present study (Table S1).

For each candidate cargo, we generated AlphaFold 3.0 models of the full-length cargo protein bound to XPO1, Ran(GTP), and RanBP1, using the amino-acid sequences corresponding to the chains in the experimentally defined structure of XPO1-Ran(GTP)-RanBP1 in complex with NES (PDB ID: 6CIT) (Figure 2A). This approach reduces the conformational search space and minimizes de novo assembly artifacts, enabling a pocket-level assessment of whether candidate segments adopt a bona fide NES pose within the H11-H12 of XPO1.

**Figure 2.**
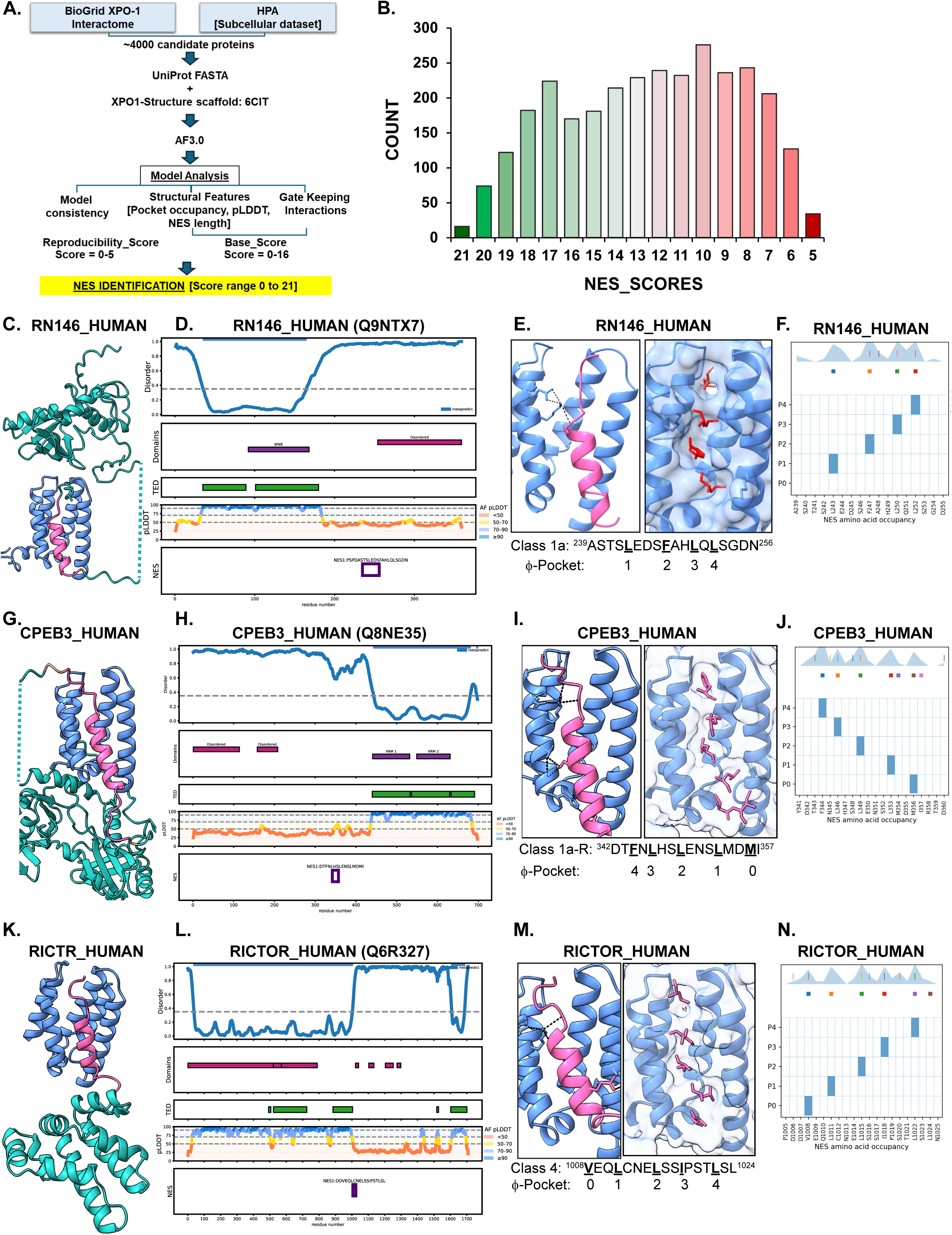
Proteome-wide, structure-resolved discovery of human NESs with AF3. **(A)** AF3-based workflow and scoring scheme for NES discovery. Candidate cargos were drawn from the BioGRID XPO1 interactome and the nucleo-cytoplasmic subset of the Human Protein Atlas (∼4000 non-redundant proteins), converted to UniProt FASTA sequences, and modeled as XPO1-RanGTP-RanBP1 export complexes using AF3 with the 6CIT scaffold. Resulting models were evaluated by a Model Analysis pipeline that integrates model-to-model consistency, structural features (pocket occupancy, pLDDT, NES length), and gatekeeping lysine interactions into a composite NES score composed of a reproducibility component (0-5) and a base structural score (0-16), yielding a 0-21 range for NES identification (yellow box). **(B)** Distribution of composite NES scores across all modeled cargos. Bars indicate the number of proteins achieving each integer NES score (5-21), with color shading emphasizing the transition from high-scoring (green) to low-scoring (red) candidates. **(C-F)** Representative Class 1a NES from RNF146_HUMAN (Q9NTX7): **(C)** AF3 complex overview with XPO1 (light-blue helix) and RNF146 (teal); the predicted NES from RNF146 (magenta) binds the H11-H12 groove in canonical forward Class 1a orientation. **(D)** NES plot locating the RN146 NES along the sequence. From top to bottom, intrinsic disorder trace (dashed line at 0.5), SMART domain annotations, structure-based TED/domain boundaries, AF pLDDT track (pLDDT Score bands at 50/70/90), and location of the predicted NES window (purple box) and the sequence. **(E)** Pocket-level NES occupancy: ribbon (left) and surface (right) representation views showing Φ-side-chain placement into P1-P4 and polar interactions (dashes) with the gate-keeper lysine K579 (human K568) and neighboring residues of XPO1. **(F)** Groove plot summarizing per-residue pocket assignment and contact density across AF3 models. The matrix (y-axis P0-P4, x-axis NES residues) marks filled squares for residues that anchor a hydrophobic pocket (Φ-pocket anchors) and small squares for hydrophobic residues (L/I/F/M/V) that contact the groove without anchoring. The shaded band above is the kernel-density estimate of cumulative side-chain contacts to the groove across models; vertical ticks indicate residues modeled in proximity to key basic/acidic groove residues (K525/K545/K579/E641). For RN146, anchors at L243→P1, F247→P2, L250→P3, and L252→P4 define a Class 1a register with no P0 anchor. **(G)** Representative Class 1a Reverse Class (1a-R) NES from CPEB3_HUMAN (Q8NE35). AF3 complex overview showing the CPEB3 cargo (teal) with its NES helix (magenta) docked in the XPO1 H11-H12 groove (light-blue surface). **(H)** NES-locus plot along CPEB3: intrinsic-disorder trace (dashed line at 0.5), domain annotations for the RRM/ZNF region and low-complexity segments, AF pLDDT track (bands at 50/70/90), and the predicted NES window (purple box) with sequence shown in the panel. **(I)** Pocket-level binding and secondary contacts: ribbon (left) and surface (right) views illustrating a Class 1a-R NES engagement across P4-P0 and polar interactions (dashes) with the gate-keeper lysine K579 (human K568) and neighboring residues of XPO1. **(J)** Groove plot (readout as in G) confirming reverse engagement of CPEB3 NES with Φ-anchors mapped to P4-P0. **(K)** AF3 complex overview showing an example of Class 4 NES, exemplified by RICTOR (teal) engaging the XPO1 H11-H12 groove (light-blue surface) via its NES helix (magenta). **(L)** NES-locus plot: disorder profile, domain/TED annotations for the scaffolding region, AF pLDDT track, and the predicted NES window (purple box) with the sequence provided in the panel. **(M)** Pocket-level binding and secondary contacts: ribbon (left) and surface (right) views showing a Class 4 arrangement with Φ-anchors placed in P1→P4 (no P0 anchor) and dashed polar contacts to K579/K568 within H11-H12. **(N)** Groove plot (interpretation as in G) displaying the canonical P1-P4 anchoring pattern for Class 4 NES.

To systematically evaluate these models, we implemented a structure-guided, unsupervised scoring framework guided by the geometric principles of NES engagement in the XPO1 groove (Figure 2A). For each model, we first identified a contiguous NES window by mapping hydrophobic side-chain insertions into the P0-P4 pockets and refining the window boundaries based on local groove contacts. We then computed a composite “base-score” from multiple structural components, including NES length, average per-residue predicted Local Distance Difference Test (pLDDT), the density of contacts with the XPO1 groove, the number of distinct pockets engaged (with a bonus score for NES helices spanning ≥3 pockets), and H-bonding to conserved support residues (Gate-keeping interactions: K525, K545, K579, K533). Per model base scores were combined with a reproducibility term, scored from 0 to 5, that measures how often a given NES window is recovered across the five AF3 models. The sum yielded a consensus NES and a total score from 0 to 21 (Figure 2B), which we used to rank candidates and prioritize structurally coherent, highly reproducible segments for manual curation (Table S2). For all scored NES, we computationally converted the structure model files into a 2D XPO1 pocket occupancy map at a single-residue resolution, which was visualized as a groove plot (Figure S2A). Groove plots provide a two-dimensional map of helix placement along the P0-P4 axis and facilitate rapid detection of forward versus reverse insertion and subsequent NES class assignment. Finally, the NES position within each cargo’s primary sequence and its structural context were visualized using a 2D Domain-NES plot that integrates domain annotations from SMART^29^ and The Encyclopedia of Domains (TED)^30^ together with the disorder predictions from MetaPredict^31^ (Figure S2B). TED catalogs structural domains from the AlphaFold database that are not captured by HMMER-based SMART, thereby incorporating TED refines and extends domain boundary annotations for the XPO1 interactome, providing a more granular view of where NES motifs are embedded within protein architecture. Together, this workflow and scoring matrix establish a standardized framework for large-scale interacting motif discovery from the modeled proteome. This approach not only recapitulates the canonical features of XPO1-dependent export but also enables unsupervised NES class assignment, filtering out spurious XPO1 contacts, and prioritization of motifs most likely to represent functional export signals. Applying these criteria, we identified NES-like sequences in 3,005 of the ∼4,000 proteins in our database, with each selected protein containing at least one scored NES candidate.

To benchmark our NES scoring system, we analyzed the distribution of maximum scores across candidate motifs. The highest attainable score of 21 reflects reproducible polar interactions, recurrence across AF3 models, and a consistent structural engagement matrix, including all five XPO1 pocket engagements through hydrophobic anchors in the NES sequence (Table S2). Many well-characterized NES scored in the upper range (15-21), consistently displaying robust pocket occupancy and stabilizing contacts. Several experimentally validated motifs, including RIOK2, achieved a maximum score of 21, whereas others, such as PDPK1, Cyclin B1, and APC NES1, scored slightly lower but still exhibited canonical helical placements within the groove. Reduced scores typically reflect variability in polar contacts, model reproducibility, or the absence of one of the pocket anchors (most often P0). Overall, scores of 10-21 corresponded to progressively higher confidence in NES detection, with values > 12 reliably identifying export-competent motifs with high reproducibility across models (Figure 2B, Table S2).

Using this scoring matrix, we found that the vast majority of high-confidence hits corresponded to previously unannotated XPO1 cargos, underscoring the ability of AF3-based modeling to expand the known repertoire of NES-bearing proteins. NES cargos were broadly distributed across key biological functions (Figure S2C). Transcription factors represented the largest group (506, 33.7%), followed by mRNA biology factors (196, 13.0%), Ser/Thr kinases (141, 9.4%), epigenetic/chromatin regulators (141, 9.4%), and cell-cycle regulators (157, 10.4%). Additional categories included protein phosphatases (99, 6.6%), E3 ubiquitin ligases (83, 5.5%), cancer-linked proteins (45, 3.0%), autophagy-associated factors (68, 4.5%), mitochondrial proteome components (19, 1.3%), RQC pathway components (19, 1.3%), centromere proteome factors (16, 1.1%), and SUMOylation-associated proteins (13, 0.9%).

This enrichment of transcription factors, kinases, epigenetic regulators, ubiquitin ligases, phosphatases, mitochondrial proteins, and autophagy regulators underscores the role of XPO1-dependent export in coordinating transcriptional programs, signaling pathways, proteostasis, and organelle communication. Together with the substantial representation of cancer-linked proteins, these findings highlight the clinical importance of systematically mapping NESs in the context of dysregulated nucleocytoplasmic transport.

To illustrate the range of NES structural geometries captured by AF3, we feature representative high-scoring NESs from three diverse NES classes: Class 1a (canonical), Class 1a-R (reverse orientation), and Class 4 (NES with internal-proline-dependent kink), which exemplify AF3’s ability to model diverse NES engagement with XPO1 observed across the proteome, for which no prior structural information was available.

RNF146 is a PAR-binding E3 ubiquitin ligase central to DNA damage-associated proteolysis^32^. AF3 modeling identified a conserved, high-scoring segment spanning residues 243-252 in the disordered region C-terminal to the WWE domain as a putative NES (Figure 2C-D). Notably, earlier work mapped XPO1-RNF146 interaction to a broader C-terminal segment that encompasses this region, but did not resolve a discrete NES within it^33^. Our AF3-predicted NES falls within this previously defined XPO1-interacting segment, providing a structural rationale for its exportin interaction and illustrating AF3’s ability to identify previously unrecognized functional NESs within experimentally mapped binding regions. This region forms an amphipathic α-helix that is deeply inserted into the XPO1 active site (Figure 2E). As visualized through the groove plot, hydrophobic anchors L^243^, F^247^, L^250^, and L^252^ occupy P1-P4 sequentially, in 3-2-1 spacing to match Class 1a geometry (Figure 2E-F). The NES is further stabilized by backbone hydrogen bonding to XPO1 K^579^ (Figure 2E).

CPEB3 (Cytoplasmic Polyadenylation Element-Binding protein 3) and EAF1 (Figure S2A) are examples of NES that engage with XPO1 in reverse orientation. CPEB3 is a neuron-enriched RNA-binding regulator with two RRM domains and a C3H1-zinc-knuckle that controls the translation and localization of synaptic mRNAs (Figure 2G-H). It shuttles between the nucleus and cytoplasm via XPO1-mediated export, thereby linking nuclear RNA handling to cytoplasmic translational control^34^. In our AF3 ternary model, a single high-confidence NES immediately N-terminal to the RRM tandem (Figure 2H, residues 342-357; DTFNLHSLENSLMDMI) forms an amphipathic α-helix that docks reverse-oriented Class-1a (1a-R) across the XPO1 groove. Hydrophobic anchors pack sequentially into the Φ-pockets (F344→P4, L346→P3, L353→P2, M356→P1, and I357→P0), creating a continuous P4-P0 engagement consistent with a fully competent export signal (Figure 2I-J). The helix is reinforced by the canonical backbone hydrogen-bonding network to the groove floor and by polar contacts with XPO1 K^579^ and K^545^, while acidic/polar side chains (e.g., D342, N345, H347, E350, and N351) remain solvent-oriented at the rim (Figure 2I).

Class 4 is a structurally distinct NES marked by a proline-dependent bend in the final gap between the P3 and P4 anchors (Φ3XXXΦ4)^35^. In RICTOR, an essential mTORC2 subunit, we identify a high-confidence Class 4 NES at residues 1006-1024 (DDVEQLCNELSSIPSTLSL; score 20). AF3 models place hydrophobics at the pocket anchors and locate the required proline within the Φ3-Φ4 gap (**I**PST**<u>L</u>**), generating the expected local kink while preserving pocket occupancy and polar stabilization. The forward-oriented helix spans P0-P4 and seats the widely spaced anchors V1008, L1011, L1015, I1018, and L1022 in a Class 4 register (2,3,2,3), with a supportive backbone contact to K579 (Figures 2K-N).

Together, these examples capture the structural diversity of canonical NESs predicted by AF3, from fully extended Class 1a helices to reverse-orientation and Class 4 arrangements with internal helical kink, highlighting the capacity of the XPO1 groove to recognize multiple binding geometries within a conserved pocket.

### AF3 models expand the structural geometry of NES occupancy in the XPO1 groove

To further capture the structural diversity of the NES-XPO1 interaction, we attempted to classify all NES peptides identified in our datasets. We first attempted to classify these NES into canonical NES classes (Classes 1a-d, forward and reverse, and 2, 3, and 4, Figure S1B) using an unsupervised approach to identify NES classes based on hydrophobic spacing and residue tolerance criteria (Table S3). However, while we correctly classified many known NES sequences, a substantial number (1652/3005, 53%) remained unclassified, including many high-scoring (>10) NES (Table S3). Inspection of groove plots of many classified and undefined NESs revealed that NES helix orientation in the XPO1 groove often did not match the class assigned from hydrophobic spacing, suggesting that sequence-based rules alone are insufficient.

To address this, we developed a structure-aware NES classification strategy (Figure 3A) that combines (i) the orientation of helix insertion into the XPO1 groove (forward versus reverse), as determined from NES groove plots (Figure S2A, 3B), with (ii) explicit mapping of hydrophobic residues (LIVFM) that occupy pockets P0-P4, from which we derive inter-anchor residue spacings. We then use this integrated orientation, pocket map, and spacing information to classify predicted NES helices according to canonical NES spacing rules (Orientation-aware-NES-classification, Figure 3C, Table S4). This approach validated many previously identified canonical NES classes using sequence-based methods, while resolving numerous cases of incongruence between sequence- and structure-based classifications of well-characterized NES motifs. For example, in the crystal structure of SMAD4-NES bound to XPO1 (PDB ID: 5UWU)^19^, the SMAD4 NES (^141^IDLSGLTLQ^149^) occupies the P1-P4 pockets in a manner consistent with Class 2 NES engagement (spacing: 1,2,1). In contrast, the AF3 model consistently suggests that within the full-length protein context, SMAD4 NES binds XPO1 in a reverse orientation, occupying pockets P4-P1 in 1,2,1 spacing, thereby reclassifying it as a reverse Class 2 NES (2-R) (Figure S3A-D, Table S4). Thus, we propose that NES orientation within XPO1—forward or reverse—is primarily dictated by the local conformational constraints of the cargo protein in its full-length context, rather than being inherently encoded in the NES sequence itself, especially given that XPO1 can accommodate the same NES peptide in both directions.

**Figure 3.**
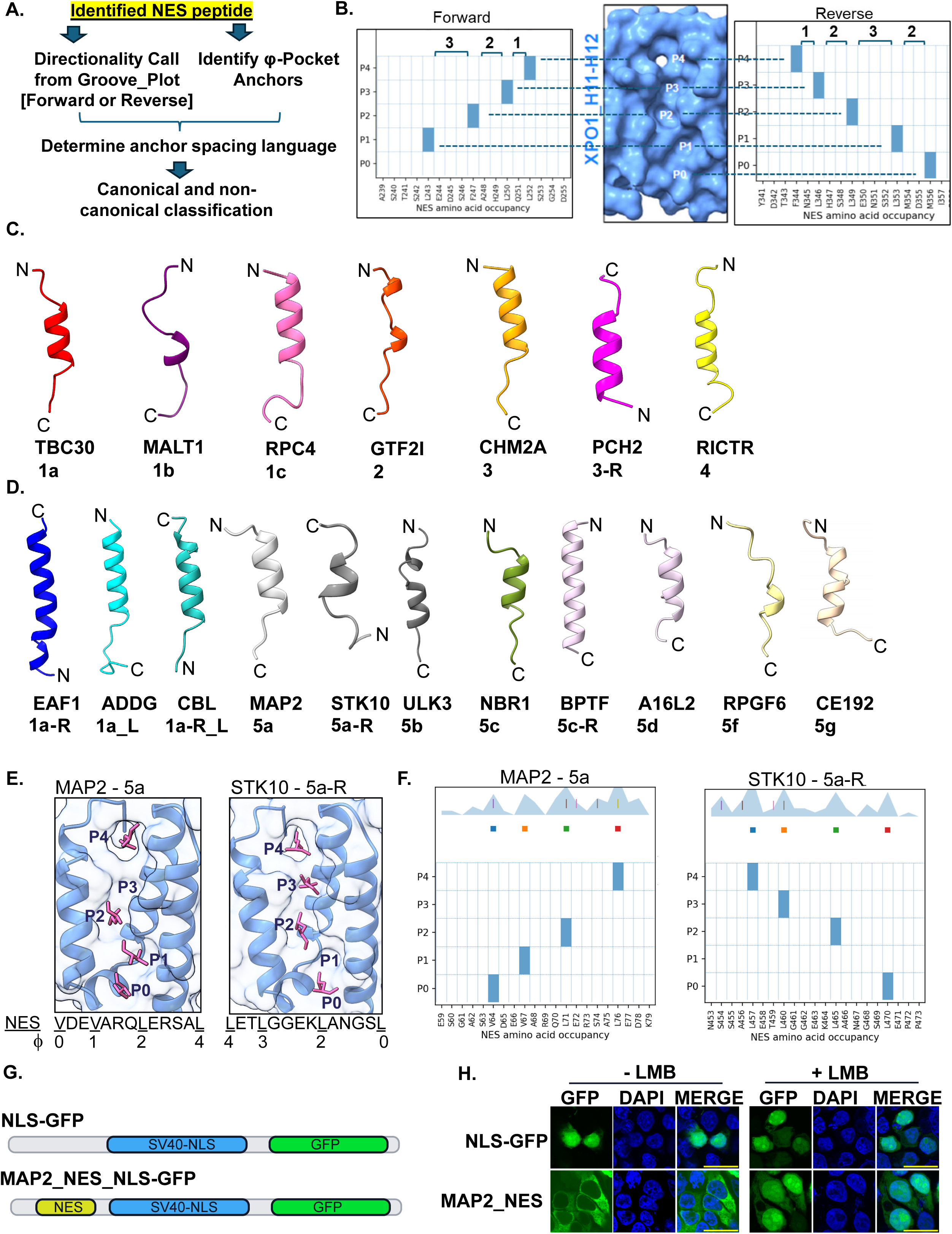
Structure-aware pipeline for NES orientation and class assignment. **(A)** Schematic of the orientation-aware, structure-aware NES classification workflow. For each identified NES peptide, groove plots are used to call directionality (forward or reverse) and to assign φ-pocket anchors (P0-P4); the resulting inter-anchor spacing language is then mapped onto canonical or non-canonical classes. **(B)** Illustration of the structure-aware classifier. Representative groove plots for forward (left) and reverse (right) NESs are shown alongside the XPO1 H11-H12 groove (center), with bars indicating pocket occupancy (P0-P4) and numbers denoting inter-anchor spacing. Together, these views demonstrate how pocket usage and orientation in the AF3 model are converted into NES class calls. **(C)** Representative NES helices spanning the canonical families. AlphaFold3 models of isolated NES segments are shown as cartoons with N- and C-termini indicated, illustrating the diversity of Φ-spacing solutions that still seat into the XPO1 groove. Examples include a canonical Class-1a NES (TBC30), Class-1b (MALT1), Class-1c (RPC4), Class-2 (GTF2I), Class-3 (CHM2A), reverse Class-3 (PCH2, 3-R), and Class-4 (RICTR). **(D)** Examples of reverse and “like” variants and the Class-5 family. AF3 models are shown for a reverse Class-1a NES (EAF1, 1a-R), a 1a-like NES (ADDG, 1a_L), a reverse 1a-like NES (CBL, 1a-R_L), and a panel of Class-5 subclasses (MAP2 5a, STK10 5a-R, ULK3 5b, NBR1 5c, BPTF 5c-R, A16L2 5d, RPGF6 5f, CE192 5g), highlighting variation in helix length, register, and amphipathic pattern while maintaining XPO1-compatible geometries. **(E)** Structural basis of Class-5 pocket usage. AF3 models of the XPO1 export complex are shown for MAP2 (Class-5a) and STK10 (Class-5a-R) NESs. The XPO1 H11-H12 groove is in blue cartoon and NES side chains occupying P0-P4 are in magenta, with pocket labels indicated. NES sequences beneath each panel mark the hydrophobic anchor positions (Φ0-Φ4) and inter-anchor gaps, illustrating the internal-skip spacing that defines Class-5a/5a-R. **(F)** Groove plots for MAP2 and STK10 NESs. Pocket-occupancy plots derived from AF3 models show the positions of residues seated in P0-P4 along the NES sequence for MAP2-5a (left) and STK10-5a-R (right), providing the orientation- and structure-aware readout used for Class-5 calling. **(G)** Design of NES reporter constructs. Schematic of the NLS-GFP control (C-terminal SV40 NLS fused to GFP) and the MAP2_NES_NLS-GFP reporter in which the MAP2 Class-5a NES is inserted N-terminal to the SV40 NLS. **(H)** Functional validation of the MAP2 NES. Fluorescence microscopy of cells expressing NLS-GFP or MAP2_NES_NLS-GFP shows nuclear localization of the control reporter and robust cytoplasmic redistribution driven by the MAP2 NES under basal conditions (-LMB). Treatment with leptomycin B (+LMB) restores nuclear GFP in the MAP2 reporter, confirming XPO1-dependent export.

Furthermore, in class IIa histone deacetylases HDAC4 and HDAC5, the C-terminal NES motifs (HDAC4: ^1048^EAETVTAMASLSVGVK^1067^ and HDAC5: ^1079^EAETVSAMALLSVGA^1096^) were annotated initially as Class 1b based solely on their hydrophobic spacing (HDAC5: E<u>A</u>ET<u>V</u>SA<u>M</u>AL<u>L</u>S<u>V</u>GA, Spacing: 2,2,2,1). However, structural work on CRM1(XPO1)-bound HDAC5 NES peptides showed that this motif actually engages the CRM1 groove, using a Class 1a pocket pattern (HDAC5: <u>E</u>AE<u>T</u>VSA<u>M</u>AL<u>L</u>S<u>V</u>GA, 2,3,2,1)^19^. Our structure-aware classifier recapitulated this geometry in AF3 models and accordingly reassigned both HDAC4 and HDAC5 NESs to Class 1a, demonstrating its ability to resolve sequence/structure incongruency (Table S4). Similarly, AF 3.0 identified the two novel NESs in the E3 ligases PJA1 and PJA2, which were labeled as Class 1b by sequence-based criteria (PJA1/PJA2:^547/586^AM<u>E</u>TA<u>L</u>AH<u>L</u>ES<u>L</u>A<u>V</u>DV^562/609^, spacing: 2,2,2,1) (Table S3), yet their pocket-resolved engagement with XPO1 matches a canonical Class 1a pocket-occupancy pattern (A<u>M</u>ET<u>A</u>LAH<u>L</u>ES<u>L</u>A<u>V</u>DV, Spacing: 2,3,2,1) (Table S4). Together, these examples underscore the importance of incorporating structural information into NES classification schemes, rather than relying solely on hydrophobic spacing patterns, especially when annotating large NES datasets.

Because our structure-aware classifier applies stricter orientation and geometric criteria and therefore prioritizes specificity over breadth, a larger fraction of NES candidates (85%) remained unclassified compared to 53% using the sequence-based classifier (Table S4). Importantly, a notable subset (Over 1000) with higher confidence scores (>10) showed reproducible pocket occupancy but with non-canonical spacing and/or non-canonical residue occupancy at one of the pockets, P0-P4. This pattern suggests the existence of additional, non-canonical NES topologies and anchor-residue usages beyond the current geometric classes and highlights these motifs as candidates for further study. Closer examination of P0-P4 pocket engagement in these high-confidence models revealed diverse anchor geometries and residue tolerances at specific pockets, extending beyond canonical allowances that allow any residue at P0 and A/T at P1 or P2, as described below:

#### Canonical-Like NES geometry

Within the group of unclassified high-confidence groove-seated NES helices, we identified a recurring subclass that preserved canonical spacing and pocket geometry but featured non-hydrophobic residues at the P3 or P4 anchor positions. For example, these motifs maintained the 3-2-1 inter-anchor spacing characteristic of Class 1a, but substituted a non-hydrophobic residue, typically threonine (T) or Alanine (A), at one of the terminal anchor positions, Φ3/Φ4 (Table S5). These motifs were designated as “Class XX_L” (e.g., 1a_Like) and “Class XX-R_L (e.g., 1a-R_L, Reverse-oriented counterparts).

One example of Class 1a_L geometry is the ADDG NES (residues 642-651: <u>L</u>AKR<u>V</u>SR<u>L</u>S<u>T</u>), which displays canonical Class 1a spacing (3-2-1) and uses well-seated hydrophobic anchors at P1-P3, but terminates with a threonine at P4. In the AF3 models, Thr^651^ adopts a rotamer in which its γ-methyl group is reproducibly facing the P4 hydrophobic pocket, occupying essentially the same groove position occupied by a hydrophobic side chain in canonical Class 1a NESs, while the hydroxyl group remains solvent-exposed (Figure S3E-F). This structural arrangement demonstrates that the P4 anchor function can, in some cases, be fulfilled by a non-hydrophobic side-chain. On this basis, we extended the classification of such motifs to “Class 1a_L” (Class 1a-like) as they align spatially and geometrically with canonical 1a registers but fail the strict hydrophobicity criterion at the P4 anchor position.

By incorporating side-chain seating directly from AF3 XPO1-NES groove models, this subclassification resolves ambiguities that arise from sequence-only rules. In our scheme, XX_L and XX-R_L variants are defined for all canonical families (Classes 1a-1d, 2, 3, and 4) by allowing alanine (A) or threonine (T) at the distal anchor positions (typically P3 and/or P4), provided that the remaining anchors follow the canonical hydrophobic pattern and the overall pocket register is preserved. In this framework, the A/T relaxation at P3-P4 in XX_L classes can be viewed as a mirror image of the established rule that alanine or threonine is tolerated at the P1-P2 anchor positions in canonical Classes 1-4. For Class 4-like motifs, we also relaxed the requirement for an intervening Pro between P3 and the P4 segment, provided the NES maintains hydrophobic anchors and overall spacing consistent with Class 4 geometry. One example is the ACBD5 NES (^164^<u>L</u>EKISKC<u>L</u>ED<u>L</u>GNV<u>L</u>^178^), in which P0-P4 are occupied by L^164^, I^167^, L^171^, L^174^, L^178^, yielding a 2-3-2-3 spacing pattern that matches the extended Class 4 register (Figure S3G-H), but without a Pro in the P3-P4 segment. AF3 models show that this C-terminal segment remains helical and that the Leu^178^ side chain is well seated in the distal P4 pocket, despite the absence of a Pro-induced turn (Figure S3G-H). Accordingly, our classifier assigns this motif as Class 4_L, reflecting canonical Class 4 spacing and pocket geometry with a relaxed requirement for a Pro-containing tail. Thus, XX_L or XX-R_L classes retain the canonical residue spacing rule, while expanding the Alanine/Threonine usage rule at distal pockets, resulting in the identification of 174 proteins with NES belonging to the canonical-Like classes in our datasets, featuring A/T at either P3 or P4 position.

#### Class 5: NES with an internal pocket skip

We noticed a set of high-scoring NES with a non-canonical topology defined by a single internal pocket containing a weakened (non-hydrophobic, non-A/T) residue anchor, followed by distal re-occupation. In contrast to terminal-skip variants seen in many canonical NES classes (e.g., P4 skip in Class 3 or P0 skip in Classes 1a/1b/1c/1d/2/4), these NESs, which now belong to Class 5, retain an amphipathic helix and feature terminal hydrophobic anchors at both P0 and P4, while tolerating one non-hydrophobic or unseated residue at a single internal pocket (P1, P2, or P3, depending on subclass). In the forward oriented pocket frame, the predominant subclasses cluster into P3-weak (P0-P1-P2-x-P4; Classes 5a/5b), P1-weak (P0-x-P2-P3-P4; Classes 5c/5d/5e/5f), and P2-weak (P0-P1-x-P3-P4; Class 5g) geometries, where x is any residue except LIVMFAT. We designate these as Class-5 NESs and describe them by inter-anchor gap tuples: 5a (2,3,4), 5b (3,2,4), 5c (6,3,2), 5d (5,2,1), 5e (6,2,3), 5f (4,2,1), and 5g (3,6,3), with “-R” appended for reverse-orientation counterparts (e.g., 5a-R) (Figure, 3D, Table S5).

Applying this rubric, we identified 74 Class-5 NES in our dataset. Representative forward examples include MAP2 (gene: MetAP2, P0, P1, P2, P4; gaps 2,3,4, Class 5a, Figure 3D-E), ULK3 (P0, P1, P2, P4; gaps 4,3,5, Class 5b), NBR1 (P0, P2, P3, P4; P1 weak/absent, Class 5c), A16L2 (P0, P2, P3, P4; gaps 6,2,0, Class 5d), RPGF6 (P0, P2, P3, P4; gaps 3,2,1, Class 5f), and CE192 (P0, P2, P3, P4; gaps 3,6,2, Class 5g) (Figure 3D). Reverse-orientation counterparts exhibit the same internal topology after mirroring, for example, STK10 (P0, P2, P3, P4; gaps 2,4,4, Class 5a-R, Figure 3E-F) and BPTF (P0, P2, P3, P4; gaps 6,3,2, Class 5c-R) (Figure 3D). These NES engagements occur reproducibly in pocket-resolved models with consistent helix seating. Class-5 NESs also retain conserved interface features characteristic of canonical NES binding, including engagement of the selectivity filter (e.g., K^579^ in the H11 loop) and backbone and side-chain hydrogen bonds along the groove rim. Preservation of these non-canonical internal pocket contacts explains the high structural confidence despite the weakened internal pocket and supports their interpretation as bona fide XPO1-binding geometries.

To determine if the non-canonical structural geometry of MAP2 can function to export cargo out of the nucleus, we generated a GFP-tagged reporter plasmid carrying SV-40NLS with or without Class 5a (MAP2) NES peptide (Figure 3G). Whereas the NLS-GFP fusion was strongly nuclear in the absence of any NES sequence (Figure 3H), fusion of Class 5a NES from MAP2 redistributed the reporter to the cytoplasm, indicating efficient export capabilities despite its non-canonical pocket-use topologies (Figure 3H). When cells were treated with the XPO1 inhibitor Leptomycin B (LMB), we observed strong nuclear retention of the MAP2-NES Peptide, confirming XPO1 dependency for the export of this novel Class 5 NES. Notably, a previous study reported that MAP2 is an XPO1 interactor and found that its NES is not predictable by canonical hydrophobic spacing rules^25^. AF3 modeling uncovered MAP2 NES as a founding member of Class 5 geometry.

Taken together, our proteome-wide analyses indicate that export competence is primarily governed by geometry and the pocket register - features not fully captured by linear hydrophobic consensus strings. In our pocket-based classification framework, occupancy and inter-pocket spacing define export competence, with one of the P1-P4 positions often residue-agnostic. In contrast, the others are more constrained to be hydrophobic. This understanding reconciles high-confidence cases missed by sequence rules and offers a more reliable basis for classification, prediction, targeted mutagenesis, and functional validation.

### AF3 modeling and cellular validation of NESs in cargos with disputed export signals

Our proteome-wide analysis provided an opportunity to resolve ambiguous NESs, for which previous experiments did not conclusively establish XPO1 dependence or functionality via point mutagenesis. Many such proteins harbor candidate motifs identified in silico but not validated in functional assays, leaving their classification unverified. Using our workflow, we identified high-scoring NESs in several disputed cargo proteins (Figure 4A). For further analysis, we focused on two representative cases, CEP43 and UBP12, which exemplify distinct categories of uncertainty: CEP43, where candidate NESs have been reported but remain unvalidated, and UBP12, where no NES has been previously annotated.

**Figure 4.**
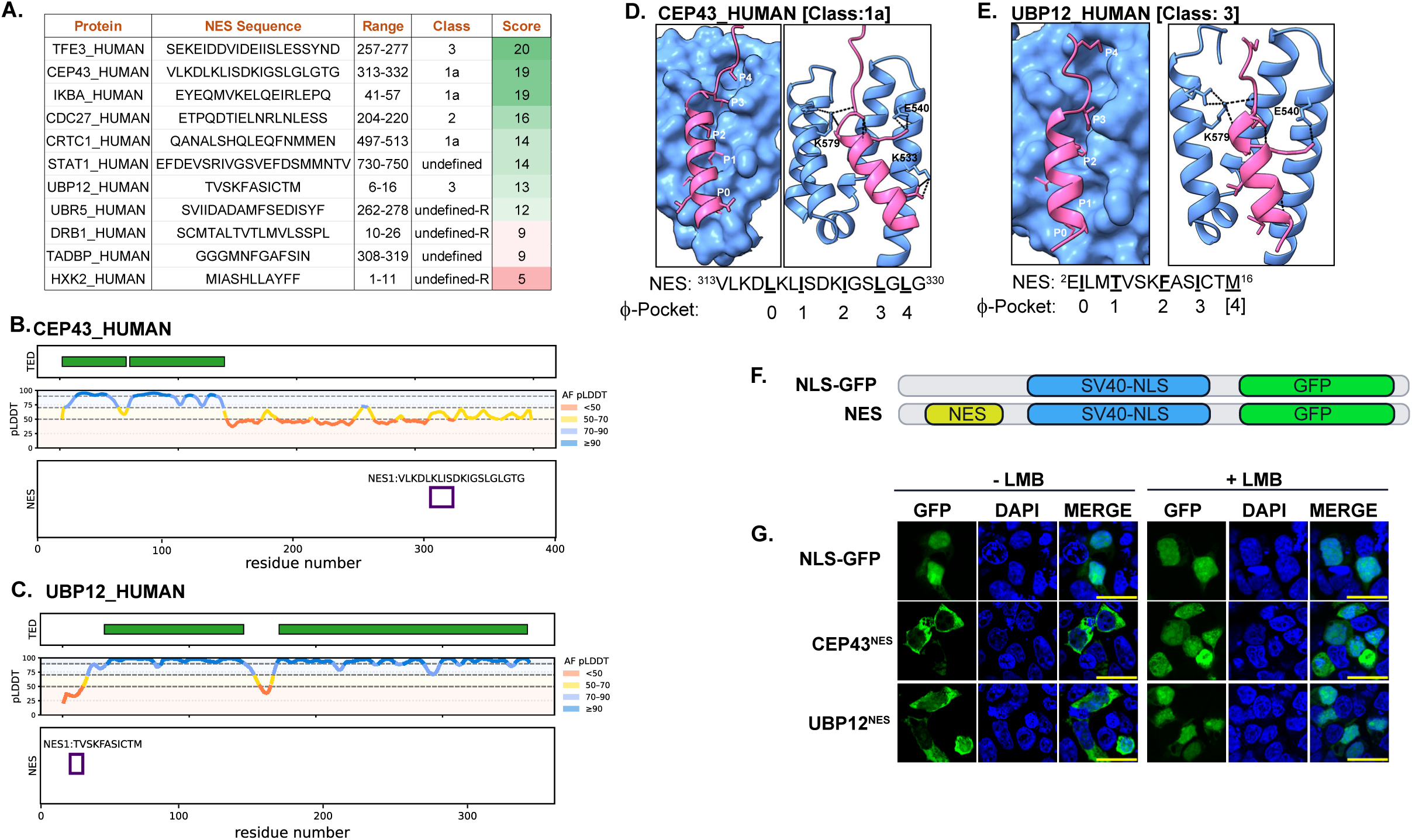
AF3 modeling and cellular validation resolve disputed XPO1 cargos. **(A)** Summary of NES sequences, positional ranges, structural classes, and composite NES scores for disputed or previously uncertain XPO1 cargos. **(B-C)** NES-locus plots for CEP43 (B) and UBP12 (C). Tracks (top to bottom) show TED/domain annotations (green bars), AF3 pLDDT along the primary sequence (color-coded at 50/70/90), and the predicted NES window (purple box) with sequence, locating each NES within the protein architecture. **(D-E)** Pocket-level AF3 export-complex models for CEP43 (D, Class 1a) and UBP12 (E, Class 3). For each cargo, left panels show XPO1 (light-blue surface) with the H11-H12 groove and the docked NES helix (magenta). Right panels show ribbon views highlighting Φ-side-chain placement into pockets P0-P4 and the polar contact network (dashed lines) involving the gatekeeper lysine (yeast K579 / human K568) and neighboring residues. **(F)** Design of NES reporters. Top, NLS-GFP control (SV40 NLS fused to GFP). Bottom, NES-NLS-GFP in which the candidate NES (yellow) is inserted N-terminal to the SV40 NLS-GFP cassette. **(G)** Cellular validation of CEP43 and UBP12 NESs. Fluorescence microscopy of cells expressing NLS-GFP, CEP43^NES^, or UBP12^NES^ reporters imaged without (−LMB) or with leptomycin B (+LMB). The high-scoring CEP43 NES (score = 19) drives robust cytoplasmic GFP accumulation under basal conditions, whereas the lower-scoring UBP12 NES (score = 13) produces comparatively weaker export with more nuclear signal. In both cases, LMB treatment restores nuclear GFP, confirming XPO1-dependent export for CEP43 (313-332) and UBP12 (6-16). Scale bars, 20 μm.

#### CEP43 (residues 313-332; NES: VLKDLKLISDKIGSLGLGTG, Class 1a)

CEP43 (FGFR1OP/FOP) is a centrosomal scaffold protein that is required for microtubule anchoring and ciliogenesis. Previous studies proposed NESs in the 352-397 region, but these displayed weak activity in Rev(1.4)-GFP assays and no XPO1 dependence in SRV_B/A reporters, leaving the NES of CEP43 unresolved^36^. AF3 consistently predicted an alternative upstream motif spanning residues 313-332 that adopts a canonical Class 1a NES configuration (Figure 4B). In the modeled complex, this segment forms a 20-residue amphipathic α-helix aligned within the HEAT 11-12 groove of NPC. Hydrophobic residues L^316^, I^319^, L^322^, L^326^, and L^330^ sequentially occupied the P0-P4 pockets, whereas D318 formed a salt bridge with K527, and the backbone carbonyls established hydrogen bonds with K^579^, anchoring the helix in a canonical orientation. These features accounted for the high structural score (19/21), consistent with high reproducibility across AF3 models.

#### UBP12 (residues 6-16; NES: TVSKFASICTM, Class 3)

UBP12 (USP12) is a deubiquitinating enzyme implicated in ubiquitin signaling and substrate turnover, but a functional NES has not been convincingly defined^37,38^. AF3 modeling consistently places residues 6 to 16 at the N terminus of UBP12 as a short amphipathic helix in a Class 3 geometry, with Thr6 at P1, Phe10 at P2, and Ile13 at P3 arranged in a Φ-x3-Φ-x2-Φ pattern within the XPO1 groove (Figure 4E). Interestingly, Ile3 at P0 and Met16 at P4 sit just outside pocket range across all models, suggesting that modest local dynamics in the cellular context could tip these edge anchors into or out of engagement and thereby tune UBP12 export. The tenuous predicted pocket-occupancy pattern at P0, P1 (T6), and P4 (M16) may explain both the relatively weak NES score (13/21) and the limited reproducibility across models (Figure 4A). Human UBP12 is a zinc-binding deubiquitinase, consistent with both UniProt annotation and the X-ray structure in which Zn²⁺ is resolved in complex with UBP12^39^. Explicit inclusion of Zn²⁺ during structural modeling improved NES-peptide binding within the XPO1 groove, increasing the score to 15/21 due to improved NES contact with XPO1 pockets (Table S2).

To validate the AF3 predictions, we tested whether the identified NESs from CEP43 and UBP12 drive XPO1-dependent nuclear export. We cloned the predicted segments from CEP43 (residues 313-332) and UBP12 (residues 6-16) into an SV40 NLS-GFP reporter (Figure 4F) and assessed their subcellular localization. Compared with the control SV40-NLS plasmid, which was exclusively nuclear, both NESs produced cytoplasmic reporter localization under basal conditions (Figure 4G). Interestingly, the high-scoring CEP43 NES (score = 19) drove substantially stronger export of the reporter than the UBP12 NES (score = 13), indicating that our scoring matrix can broadly distinguish potent from weaker NES elements. Treatment with leptomycin B (LMB) abolished export and redirected GFP to the nucleus for both reporters, confirming XPO1 dependence (Figure 4G). Together, these results validate the AF3-guided, structure-based discovery of functional NESs in CEP43 and UBP12 and resolve previous ambiguity surrounding their NES assignment.

### Macroautophagy Pathway Is Enriched for NES Motifs

In our proteome-wide discovery dataset, the macroautophagy pathway emerged as a prominent hotspot for putative nuclear export signals (NESs) (Figure 5A). High-confidence predictions spanned all major stages of autophagosome biogenesis and trafficking, including initiation (ULK/ATG1 complex), conjugation (ATG12-ATG5-ATG16 axis and associated E1/E2/E3-like enzymes), membrane expansion (ATG2 paralogs), and vesicle transport/maturation (FYCO1, UVRAG, and AMBRA1). This widespread distribution suggests that nucleocytoplasmic transport is integrated into autophagy regulation at multiple nodes rather than being confined to a single control point.

**Figure 5.**
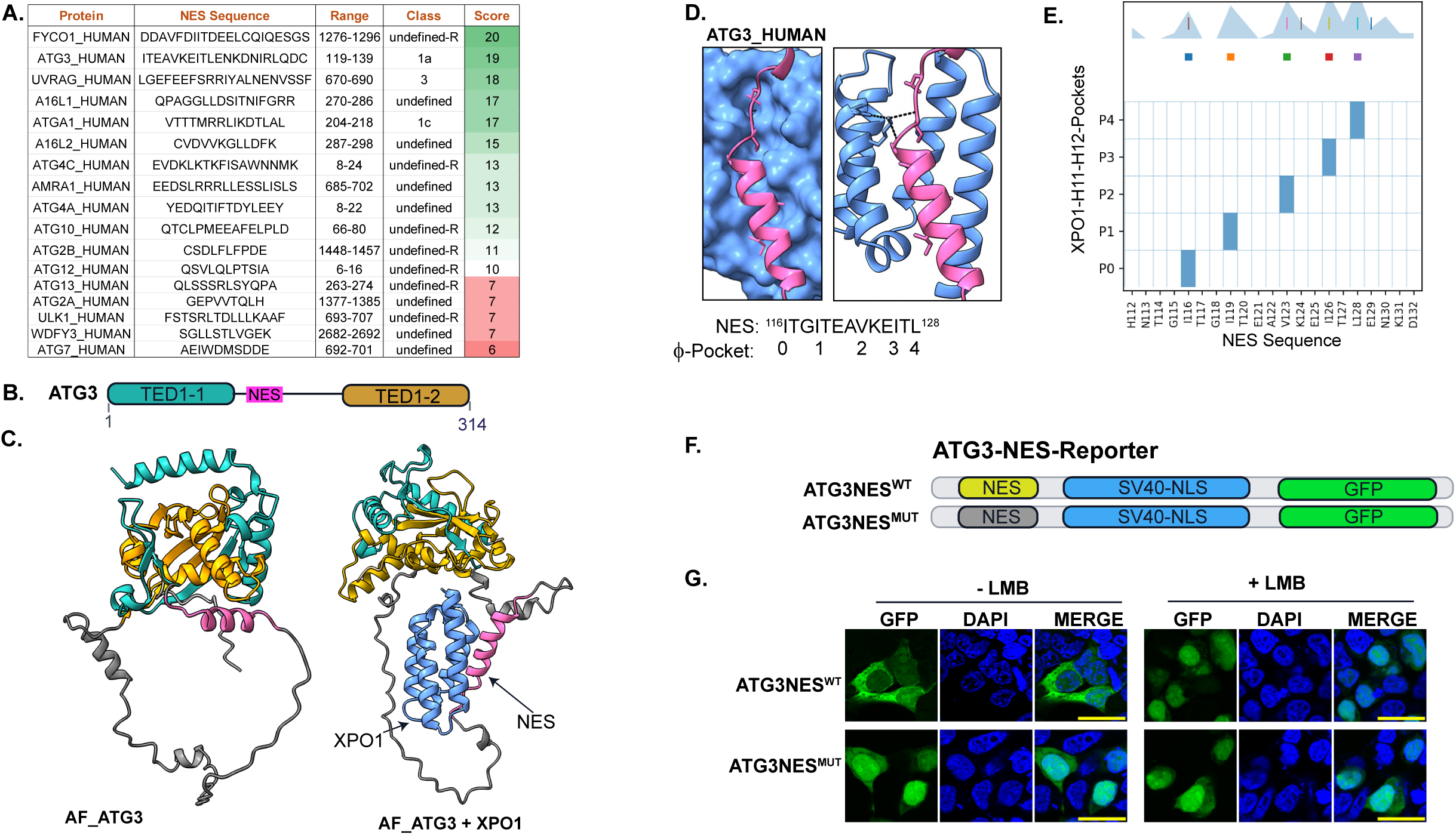
Identification and validation of an XPO1-dependent NES in ATG3 among macroautophagy components. (A) High-scoring NES calls in core macroautophagy proteins, ranked by composite NES score; ATG3 is highlighted as a top-scoring candidate (score = 19). (B) Linear domain map of human ATG3 showing the TED1-1 and TED1-2 domains flanking an inter-domain linker that contains the predicted NES. (C) AF3 models of ATG3 alone (left) and in complex with XPO1 (right). In the apo model, the linker/NES (magenta) is partially occluded between TED1-1 (cyan) and TED1-2 (orange), whereas in the export complex, the linker forms a stable helix that is fully exposed and docked into the XPO1 H11-H12 groove. (D) Pocket-level view of the AF3 ATG3-XPO1 complex. XPO1 is shown as a light-blue surface or ribbon and the ATG3 NES helix in magenta, with Φ side chains seated in hydrophobic pockets P0-P4 and polar contacts (dashed lines) to the gatekeeper lysine K579 (human K568). (E) Orientation- and structure-aware groove plot for the ATG3 NES, reporting per-residue occupancy of pockets P0-P4 and illustrating a Class-1a spacing pattern. (F) Design of ATG3 NES reporters. ATG3NES^WT^ and ATG3NES^MUT^ (hydrophobic anchor residues mutated to disrupt pocket binding) are fused N-terminal to an SV40 NLS-GFP cassette. (G) Cellular validation of the ATG3 NES. Fluorescence microscopy of cells expressing ATG3NES^WT^ or ATG3NES^MUT^ reporters treated with vehicle (-LMB) or leptomycin B (+LMB). ATG3NES^WT^ drives robust cytoplasmic GFP localization that collapses to the nucleus upon LMB treatment, whereas ATG3NES^MUT^ remains predominantly nuclear under both conditions, confirming an XPO1-dependent NES in ATG3. Scale bars, 20 μm.

Several candidates scored in the top tier of our structural metrics and were consistently reproduced across the AF3 model indices. These include ATG16L1 (Q676U5; residues 275-286; sequence: LLDSITNIFGRR), ATG3 (Q9NT62; 119-138; ITEAVKEITLENKDNIRLQD), FYCO1 (Q9BQS8; 1276-1295; DDAVFDIITDEELCQIQESG), and UVRAG (Q9P2Y5; 672-690; EFEEFSRRIYALNENVSSF). These motifs exhibited robust groove engagement and satisfied core structural criteria, including a hydrophobic anchor registry across XPO1 pockets P0-P4 and backbone contact patterns consistent with Lys^579^. Many high-scoring NES for proteins involved in macroautophagy use non-canonical anchor spacings and thus remain unclassified. These include NES for FYCO1 (Score = 20), A16L1 (Score = 17), and others (Figure 5A).

Previous reports have identified ATG3 as an interactor of XPO1 using high-throughput interactome screening methods^25^. ATG3 is an E2-like conjugating enzyme essential for autophagy. It catalyzes the lipidation of LC3/ATG8 through a thioester intermediate using its catalytic cysteine. Structurally, ATG3 contains an atypical E2 fold interrupted by large insertions that contribute to partner interaction and membrane binding (Figure 5B). Despite proteomic evidence showing the interaction between ATG3 and XPO1/XPO1 and the leptomycin B (LMB)-dependent nuclear accumulation of ATG3, its NES motif has not yet been defined.

Based on the full-length AF2.0 structure of ATG3, TED-based domain mapping partitions the protein into two subdomains, TED1-1 (residues 1-118) and TED1-2 (residues 139-314) (Figure 5B-C). AF3 identified a high confidence NES (residues 116-128, ITGITEAVKEITL, Score 20) within the linker that connects these split catalytic subdomains, which together form a composite E2 core. In models generated without XPO1, ATG3 adopts a compact conformation in which this linker is partially buried, and the NES remains disordered and occluded by intramolecular contacts between TED1-1 and TED1-2 (Figure 5C). In contrast, in ATG3-XPO1-RanGTP-RanBP1 complexes, AF3 predicts a pronounced rearrangement where TED1-1 and TED1-2 rotate to expose the linker, and the NES forms a stable amphipathic alpha helix that docks into the H11-H12 groove of XPO1 (Figure 5C-E). Sequence-based spacing and structure-aware classifier converge on Class 1a (^116^ITGITEA<u>V</u>KEIT<u>L</u>^128^, with 2-3-2-1 spacing) geometry for this NES.

To test whether the AF3-modeled NES is functional in cells, we fused the ATG3 NES sequence to an SV40 NLS-GFP reporter (Figure 5G, NES-SV40-NLS-GFP; ATGNES^WT^). In mammalian cells, the ATG3NES^WT^ efficiently overrode the strong SV40 NLS, driving reporter accumulation in the cytoplasm, consistent with active nuclear export. Treatment with leptomycin B (LMB) resulted in nuclear retention of the ATG3 reporter, demonstrating XPO1 dependency for nuclear export. In contrast, a NES-mutant reporter in which hydrophobic anchors predicted to occupy P1/P3/P4 were mutated (I119S, I126S, and L128S) was exclusively nuclear, regardless of leptomycin B (LMB) treatment. Together, these data confirm that the AF3-predicted motif is a bona fide XPO1-dependent NES in ATG3, capable of exporting NLS-tagged cargo from cells (Figure 5G).

### Identification and validation of NESs in the Ribosome Quality Control (RQC) pathway

In our proteome-wide discovery dataset, the ribosome quality control (RQC) pathway emerged as a surprising hotspot for high-scoring nuclear export signals (NESs) (Figure 6A). Among the top-ranked proteins, RQC components, including ZNF598, NEMF, UBXN7, UFD1, TERA, and ANKZ1, were consistently recovered with the canonical NES helices. Together, these proteins span nearly every functional node of the RQC cascade, from collision sensing and stalled-60S processing to ubiquitylation, extraction, and peptidyl-tRNA hydrolysis^40,41^. As RQC has traditionally been viewed as a cytoplasmic and ER-proximal process with no defined nuclear steps, the unexpected clustering of XPO1-compatible NESs in this pathway highlights a potentially overlooked nuclear interface for translation-coupled quality control. Therefore, we analyzed XPO1-NES structure for each component of the RQC pathway in detail to define the potential implications of XPO1-dependent regulation.

**Figure 6.**
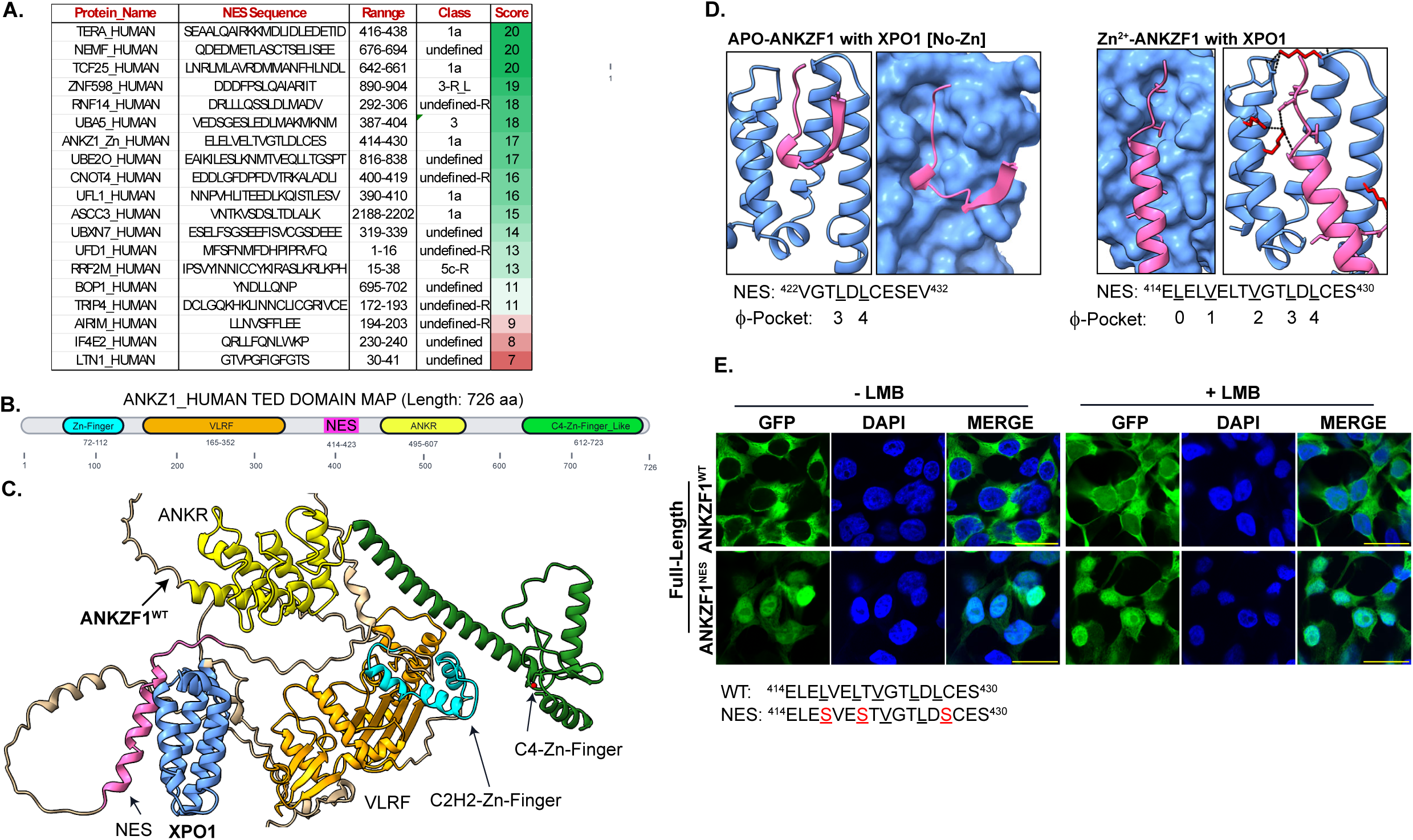
ANKZF1 harbors an allosterically controlled NES within the ribosome quality-control pathway. (A) High-scoring NES calls in RQC and RQC-linked factors ranked by composite NES score; top hits include TERA/VCP, NEMF, TCF25, ZNF598, RNF14, UBA5, ANKZF1, and UBE2O. (B) Linear TED-based domain map of ANKZF1 (726 aa) showing the N-terminal C2H2 Zn-finger (72-112), VLRF domain (165-352), NES-containing linker (414-423), ankyrin repeats (493-607), and C-terminal C4 Zn-finger-like module (612-723). (C) AF3 model of full-length ANKZF1 (colored by domain) bound to XPO1 (light blue), highlighting the NES helix (magenta) positioned between the VLRF and ankyrin repeat regions and presented to the H11-H12 groove. (D) Zinc-dependent control of NES presentation. Left, apo-ANKZF1 modeled with XPO1 shows residues 422-432 forming a short, poorly engaged segment that occupies only distal pockets (P3-P4). Right, Zn²⁺-bound ANKZF1 modeled with XPO1 forms a continuous NES helix (^414^ELELELVELTVGTLDLCES^430^) that docks along the groove with Φ side chains seated in pockets P0-P4 and polar contacts (dashed lines) to the gatekeeper lysine K579 (human K568). (E) Functional validation of the ANKZF1 NES in cells. GFP-tagged full-length ANKZF1^WT^ or NES-mutant ANKZF1^NES^ (hydrophobic anchors mutated to serine, red) were expressed in HEK293T cells and imaged without (−LMB) or with leptomycin B (+LMB). ANKZF1^WT^ shows predominantly cytoplasmic localization that shifts to the nucleus upon LMB treatment, whereas ANKZF1^NES^ is constitutively nuclear and insensitive to LMB, confirming an XPO1-dependent NES in ANKZF1. Scale bars, 20 μm.

AF3 identifies export-competent NES helices across the ribosome-collision and RQC network. In the collision-sensing arm, ZNF598 harbors high-confidence, forward Class 3-R_Like helices with sequential pocket occupancy and stabilizing XPO1 contacts (e.g., K527/K579). For ZNF598, adding or omitting Zn during modeling had little effect on the local NES helix, indicating that Zn coordination does not reshape the groove-facing amphipathic surface. Within stalled-60S recognition and RQC assembly, strong NESs occur in NEMF and UBE2O, as well as in the p97/VCP axis (UBXN7, UFD1, and VCP/TERA), consistent with XPO1 control at the level of complex assembly and substrate extraction. AF3 also recovers NESs in related proteostasis pathways, including UBA5 and UFL1 in the UFMylation arm near ER-linked ribosomes, as well as CNOT4 and ASCC3, suggesting coordination between RNA decay, transcription-coupled repair, and RQC. Notably, ANKZF1’s NES is observed in the Zn-bound model but not in the absence of Zn (see below), indicating metal-dependent control of its NES exposure. Altogether, the widespread presence of high-scoring NESs across RQC components, including ZNF598, NEMF, UBE2O, RNF14, TERA, UFD1, UBXN7, and ANKZ1, supports the existence of a genuine RQC-XPO1 axis with unexplored nuclear-level roles in regulating RQC components.

### ANKZF1 is a nucleo-cytoplasmic shuttling protein that contains a cofactor-controlled nuclear export signal (NES)

To test whether components of the RQC pathway shuttle between the nucleus and cytoplasm, we focused on ANKZF1 (Vms1), an RQC factor in the ubiquitin-proteasome system. ANKZF1 is a dual-function protein that localizes to the mitochondria, where it contributes to organellar quality control^42,43^ and operates in the ribosome quality control (RQC) pathway as a peptidyl-tRNA hydrolase responsible for releasing stalled nascent chains from obstructed ribosomes^44^. Through this activity, ANKZF1 terminates aberrant translation and prevents the accumulation of incomplete proteins, serving as the terminal “release factor” in RQC.

Domain annotation of ANKZF1 based on UniProt identifies a canonical C2H2 zinc finger (residues 72-96), a VLRF domain (203-346) implicated in ribosome binding, ankyrin repeats (493-563) mediating protein-protein interactions, and a VCP/p97-interacting motif (VIM; residues 654-666) essential for engagement with the segregase complex. AlphaFold-structure guided TED domain mapping refined this annotation by extending the N-terminal zinc finger to residues 72-112, identifying a previously unannotated C-terminal Zn-finger-like module spanning residues 612-723, and delineating more sharply defined linker regions than those previously reported (Figure 6B-C).

Initial AF3 modeling of ANKZF1 in the absence of zinc (APO-ANKZF1) failed to yield a high-confidence structure in complex with the nuclear export receptor XPO1. In 2 of the 5 models, residues 422-432 partially occupied the XPO1 pocket (Figure 6D). Given the zinc-binding nature of ANKZF1, we repeated AF3 modeling with zinc coordinated to the N-terminal and/or C-terminal zinc finger domains. Under these conditions, the region occupying the XPO1 groove (^411^FQVELELVELTVGTLDLCES^430^) exhibited increased structural flexibility, high model reproducibility, improved local confidence scores, and enhanced distal Φ-pocket occupancy (Figure 6C-D). Thus, Zinc coordination promoted the tighter packing of hydrophobic residues (L415, L418, V422, V425, and L427) into the XPO1 groove and stabilized the NES helix through key intramolecular interactions, including an E414-K527 salt bridge and K579-mediated hydrogen bonding along the NES helix (Figure 6D). The ANKZF1 NES peptide engages with the XPO1 groove in Class 1a geometry, with canonical 2,3,2,1 (^415^<u>L</u>EL<u>V</u>ELT<u>V</u>GT<u>L</u>D<u>L</u>^428^) spacing along the hydrophobic pockets of XPO1 (Figure 6D).

To validate the functional relevance of the predicted NES in the full-length protein context, we assessed the subcellular localization of GFP-tagged full-length wild-type (WT) or NES-mutated ANKZF1 proteins (Figure 6E). Wild-type ANKZF1 was predominantly cytoplasmic, but treatment with Leptomycin B (LMB), an inhibitor of XPO1-mediated nuclear export, resulted in modest but consistent nuclear accumulation, indicating that ANKZF1 is a nucleocytoplasmic shuttling protein that relies on XPO1 for nuclear export.

Consistent with this, mutations targeting hydrophobic residues critical for XPO1 groove engagement (P1, P3, and P4 pockets) abolished nucleocytoplasmic shuttling, leading to persistent nuclear retention of ANKZF1, which was not further enhanced by LMB treatment (Figure 6E).

These findings confirm ANKZF1 as an XPO1-dependent shuttling protein and demonstrate the utility of AF3 modeling in identifying previously unrecognized NES motifs. Moreover, the data highlight a cofactor-dependent mechanism by which zinc can modulate NES accessibility, suggesting a regulatory role for ANKZF1 zinc fingers in nucleocytoplasmic partitioning.

### Juxtaposed NES-NLS Architectures Suggest Coordinated Regulation of Nucleocytoplasmic Shuttling

While mapping the ANKZF1 NES (residues 414-426), we observed adjacent clusters of basic residues resembling canonical monopartite nuclear localization signal (NLS) architectures spanning residues 434-446 immediately C-terminal to the NES (Figure 7A). Modeling NLS-Importin Alpha through AF3 revealed that when the NLS peptide of ANKZF1 engages importin-α, the adjacent NES becomes unstructured and solvent-exposed, lacking the defined amphipathic helix observed in the XPO1-bound configuration (Figure 7B). This suggests a mutually exclusive binding mechanism whereby in the importin-bound state, the NES is conformationally inactive for export. Such a model implies that ANKZF1 toggles between import- and export-competent conformations, perhaps in a regulated manner, thereby coupling nuclear entry and exit through a single, structurally integrated NES-NLS module.

**Figure 7.**
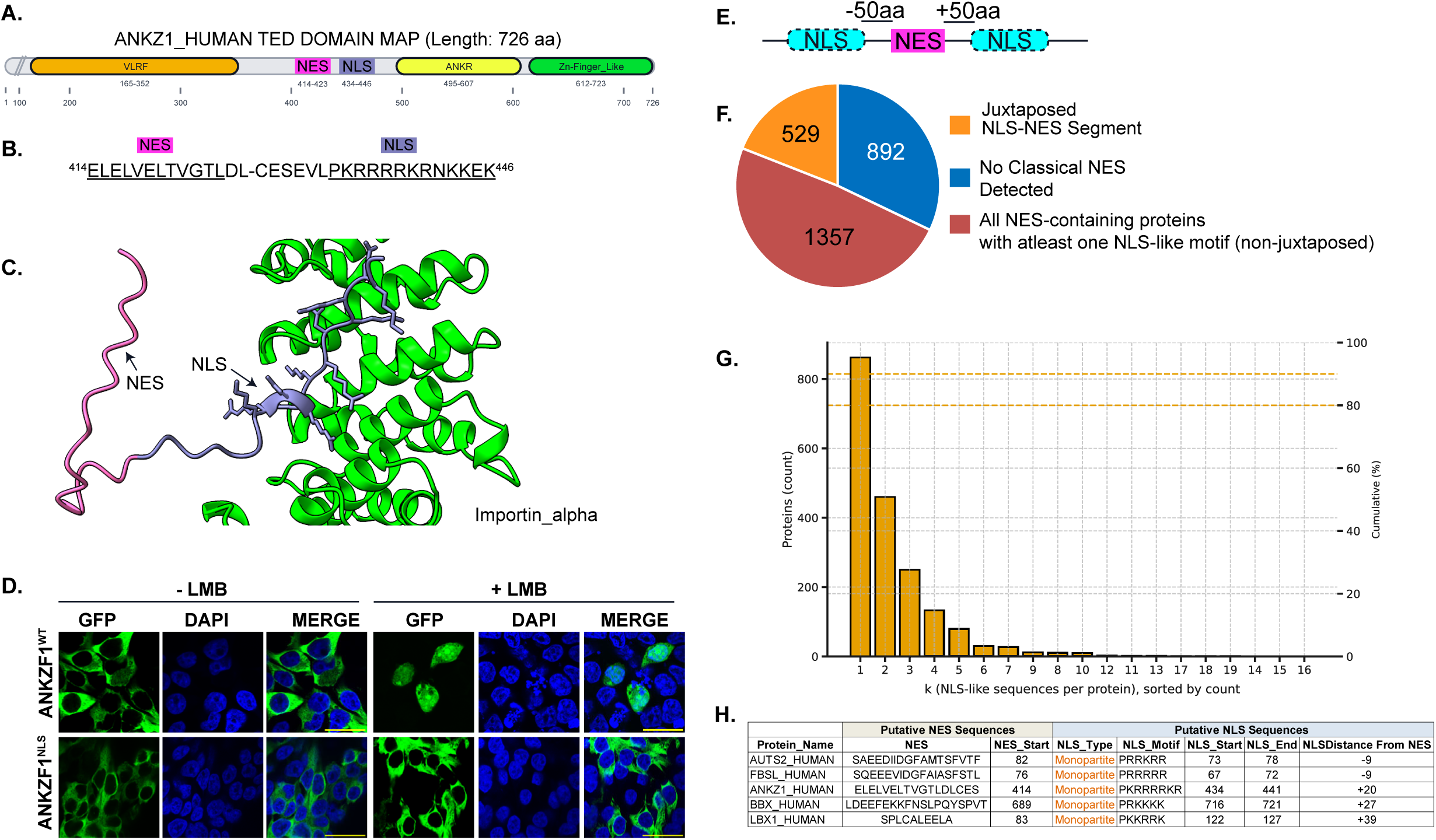
Juxtaposed NES-NLS architectures suggest co-regulation of nucleocytoplasmic shuttling across the human exportome. (A) TED-based domain map of ANKZF1 (726 aa) showing the NES (residues ∼414-423) positioned immediately adjacent to a predicted monopartite NLS (residues 434-446) within the linker between the VLRF and ankyrin-repeat regions. (B) Sequence context of the ANKZF1 module with the NES (magenta) and NLS (cyan) highlighted. (C) AF3 model of importin-α (green) bound to the ANKZF1 NLS (cyan). The adjacent NES segment (magenta) appears disordered, and solvent exposed in the importin complex, whereas in XPO1 models the same segment forms a helix that docks in the groove, consistent with mutually exclusive binding by importin and exportin. (D) Experimental validation of the ANKZF1 NLS. HEK293T cells expressing GFP-tagged full-length ANKZF1^WT^ or an NLS mutant (substitutions within residues 434-446) were treated with vehicle or 50 nM leptomycin B (LMB) for 2.5 h, fixed, stained with DAPI, and imaged by confocal microscopy under identical acquisition settings. Scale bar, 20 μm. (E) Schematic of the proteome-wide search for juxtaposed modules. Classical monopartite or bipartite NLS motifs were queried within ±50 amino acids of each AF3-identified NES. (F) Summary of outcomes for all NES-containing proteins: juxtaposed NES-NLS segments, proteins with no classical NLS detected, and proteins that contain at least one NLS but not in a juxtaposed arrangement. Counts are shown in the pie chart. (G) Distribution of the number of NLS-like sequences per NES-containing protein. Bars show protein counts for each value of k (NLS-like motifs per protein), with dashed lines indicating cumulative percentages on the right axis. (H) Representative examples of juxtaposed modules listing putative NES sequences and adjacent NLS sequences, including motif type, start and end positions, and distance from the NES (positive values, C-terminal; negative values, N-terminal).

To test whether predicted NLS plays a role in ANKZF1 nucleocytoplasmic shuttling, we generated WT or NLS-mutated [R^436^-^439^A) versions of full-length ANKZF1 fused with GFP. As expected, under basal conditions, wild-type ANKZF1 localized predominantly to the cytoplasm, due to the presence of strong NES as determined by its nuclear retention upon leptomycin B (LMB) treatment (Figure 7C). In contrast, the NLS mutant failed to accumulate in the nucleus after LMB treatment, remaining diffusely cytoplasmic (Figure 7C). These results confirm the functional role of the predicted NLS for nuclear entry and that ANKZF1 is an XPO1-dependent shuttling protein.

To assess whether tandem NES-NLS modules are broadly represented across the exportome, we scanned all high-confidence NES cargos (n = 2,778) for a classical mono- or bipartite NLS within ±50 residues of the AF3-predicted NES (Table S6, Figure 7D). This analysis identified 529 cargos (19.0%) with an NLS in this ±50-residue window; 1,357 additional cargos contained an NLS elsewhere in the protein, and 892 lacked a classical NLS (Figure 7F). These data indicate that juxtaposed import-export configurations are a common, potentially conserved feature of the human exportome.

Among proteins with at least one classical NLS (including juxtaposed sites), we quantified the number of NLS-like motifs per NES-containing cargo. The distribution was strongly right-skewed: most proteins carried one or two motifs, and the cumulative curves showed that ≈80% have ≤2 and ≈90% have ≤3 (Figure 7G). Only a small minority contained larger numbers, producing a long tail up to ∼16 motifs per protein, which likely reflects degenerate matches or context-dependent elements rather than many independent, functional NLSs.

Focusing on juxtaposed cases (NLS within ±50 residues of the AF3-predicted NES), we highlight representative NES-NLS pairs with spacing and orientation similar to the validated ANKZF1 NLS. We observed ANKZF1-like NLSs immediately upstream (e.g., −9 aa in AUTS2 and FBSL) and downstream (e.g., +27 in BBX, +39 in LBX1) of high-confidence NESs, with typical separations of ∼9-40 residues (Figure 7H). These examples reinforce a co-regulatory NES-NLS architecture in which import and export modules are arranged in tandem on the same segment, enabling spatial control of nucleocytoplasmic shuttling and providing a tractable context for regulation by local sequence changes or post-translational modifications.

Together, this study establishes a structure-resolved framework for defining export motifs across the human proteome and reveals that nuclear export functions within a bidirectional, interdependent transport system. By combining deep-learning-based structural modeling with cellular validation, we demonstrated that short peptide motifs can encode complex regulatory behaviors, bridging sequence, structure, and cellular localization. The resulting structural exportome map provides a blueprint for future studies linking nucleocytoplasmic dynamics to cell fate, signaling, and disease.

## Discussion

This study establishes a structure-informed framework for systematically identifying nuclear export signals (NESs) across the human proteome using AlphaFold 3.0 (AF3) modeling of the XPO1-RanGTP-cargo complex. Although XPO1-mediated transport has been studied extensively, the degeneracy, context dependence, and intrinsic disorder of NES motifs have hindered sequence-based identification and left many bona fide cargos undefined in the literature. By modeling over 4,000 candidate proteins in the receptor-bound state, we demonstrated that deep-learning-based structural prediction can accurately capture the geometry, pocket occupancy, and orientation of NES helices within the H11-H12 groove of XPO1. AF3 not only reproduces experimentally validated NES conformations but also uncovers previously unrecognized motifs that satisfy the biochemical and structural criteria for receptor recognition. These results transform NES discovery from a sequence motif-based search into a structure-resolved, pocket-level framework that links motif architecture and function.

### Structure-based discovery expands the human exportome

Our AF3-guided analysis identified nearly 3,000 high-confidence NESs across the human proteome, encompassing canonical and reverse orientation classes, as well as hundreds of non-canonical motifs that nonetheless exhibited stable pocket occupancy and consistent hydrophobic anchor registry. This structure-based classification refines prior large-scale XPO1 interactome studies by defining the precise physical geometry of NES-receptor engagement rather than inferring binding from co-purification data^25^. The resulting proteome-wide, pocket-resolved exportome provides a foundation for understanding how local folding, partner binding, and post-translational modifications govern NES accessibility and export competence.

### Flexibility and Regulation in XPO1-Mediated Nuclear Export

Deep learning-based AF3 modeling, combined with our statistically powerful, structure-aware NES identification framework that integrates orientation and pocket occupancy, reveals that the XPO1 groove accommodates a broader spectrum of binding geometries than previously appreciated. Canonical classes remain at the core of this family, but their “like” subclasses (for example, Class 1a-like) extend the residue-spacing language by tolerating alanine or threonine at non-canonical pockets (P3 and P4) while preserving the overall topological logic of export. Within this expanded landscape, Class 5 introduces a distinct internal-skip geometry, in which an amphipathic helix temporarily bypasses one pocket and then re-engages the groove downstream while maintaining the polar contact network, as exemplified by the MAP2 NES. Together, the canonical classes, their like counterparts, and the internal-skip arrangements demonstrate that pocket usage can be rewired without breaking the fundamental logic of nuclear export, and they illustrate how AF3 can expose subtle but reproducible alternatives that are largely invisible to sequence-based analyses alone.

Beyond these defined classes, we observed numerous high-scoring NESs that do not fit neatly into the current “Like” subclasses or the Class 5 spacing rules. For example, Cyclin B1, TRAF7, Cyclin B2, DAPK2, and COPZ1 all use a robust 3-3-0 spacing pattern that still seats into the groove but falls outside canonical Class 1a parameters. Likewise, NES motifs in PHF6 and RNF14 exhibit non-canonical pocket usage and spacing, suggesting additional tiers of internal skips or distal pocket engagement that we did not formalize in the Results section of this manuscript. We therefore restricted our primary classification to canonical, canonical_Like, and Class 5 categories, but view Cyclin B1, PHF6, RNF14, and related high-scoring motifs as candidate exemplars of higher-order, non-canonical architectures for further exploration. These cases underscore that NES space is best understood as a continuum of related geometries rather than a small set of rigid classes, and they highlight how AF3-guided modeling can uncover new, testable hypotheses about the structural grammar of nuclear export.

Flexibility in pocket occupancy by non-hydrophobic residues adds a regulatory layer to NES detection and interpretation. We repeatedly observe serine, threonine, and arginine at anchor positions, supported by local hydrogen bonds. These residues serve as substrates for post-translational modification, thereby creating natural regulatory switches. For example, in the ADDG NES, Thr^651^ occupies the P4 pocket and has been reported as phosphorylated in multiple high-throughput studies. Additionally, MAP2 contains a Class 5 NES in which S^74^ occupies the P3 pocket and is frequently phosphorylated. Whereas phosphorylation between anchors is well documented, our analysis indicates that non-canonical NESs use PTM-sensitive residues at pocket anchors themselves, enabling precise and rapid control of export activity.

Human variation and disease-linked mutations offer a second path to modulation. Substitutions that convert a serine or threonine to leucine or isoleucine at an anchor position increase hydrophobic surface area and can strengthen pocket occupancy. Structurally, such changes can transform a marginal anchor into a stable one, move a motif from an ambiguous configuration toward a canonical binding geometry, or rescue occupancy after an internal skip. This explains how conservative missense variants can shift nuclear export rates and influence dosage-sensitive transcriptional programs.

Our analysis also suggests the possibility of AF3 modeling extinct NES sequences that otherwise meet structural criteria. In several proteins, AF3 places short amphipathic segments in the groove with convincing geometry and inter-model reproducibility, even when only two or three anchors engage in the pocket. These segments could be AlphaFold “hallucinations”, but may also be vestiges that once supported export but lost function through small drifts in spacing or polarity. Even if dormant, the high-scoring models could be informative as they may mark the evolutionary sequence context of protein nuclear export. It is possible that minimal mutational events could restore function to these NES sequences, resulting in a gain-of-function.

Taken together, AF3 modeling and our structure-aware framework show that XPO1 recognizes a set of related helices that includes canonical motifs, their like subclasses with permissive spacings, and internal skip Class 5 geometries. Non hydrophobic anchors provide footholds for post translational regulation, and common clinical variants can strengthen or weaken export through simple polarity changes at key sites. Extinct NES states mapped by AF3 illuminate both evolutionary drift and practical routes to functional rescue. This view points to a next phase of experiments that pair modeling with systematic mutagenesis, including phospho mimic and methyl mimic designs, to quantify how specific chemical states move an NES along the spectrum from weak to strong export competence.

### Coupling of nuclear import and export through juxtaposed motifs

Detailed structural mapping of ANKZF1 revealed juxtaposed NLS sequences directly adjacent to their NES motifs, suggesting a design for coordinated import-export regulation in these proteins. AF3 models of ANKZF1 indicate that importin-α binding destabilizes the NES helix, rendering it inactive, whereas XPO1 binding restores helicity and export competence, a reciprocal coupling mechanism. As putative NES motifs identified in this study are validated through functional analysis, the juxtaposition of the NES-NLS sequences might illuminate the structural code of the nucleocytoplasmic shuttling machinery.

In conclusion, the structure-resolved exportome generated here provides a platform for the mechanistic annotation of pathogenic variants, offering a direct means to link sequence alterations to mislocalization phenotypes. This framework also enables the predictive testing of XPO1 inhibitors (e.g., selinexor-class drugs) in contexts where export dependence or NES integrity correlates with disease progression, thereby connecting nuclear transport modulation to therapeutic intervention.

## Methods

### Database construction

We assembled a reference table of human proteins anchored to UniProt canonical isoforms (release 2025_04), harmonizing gene names to HGNC symbols and retaining the canonical FASTA sequence plus basic metadata (length, UniProt keywords, and curated subcellular localization). Putative XPO1 cargos were compiled from human XPO1 interaction partners in BioGRID and proteins annotated as localized to both nucleus and cytoplasm in the Human Protein Atlas. After mapping to UniProt, merging, and de-duplication, and adding a small set of literature-validated NES exemplars as positive controls, we obtained a non-redundant list of ∼4,000 candidate XPO1 cargos for AF3 modeling.

### AlphaFold 3.0 modeling

Each candidate cargo was modeled as a ternary complex with human Exportin-1 (XPO1/CRM1) and Ran in a GTP-bound sequence state using the AlphaFold 3.0 (AF3) complex pipeline. Template guidance on the cargo was disabled to avoid biasing its placement in the groove, and five models (Model_0-Model_4) were generated per cargo using the auto-seed function. Per-protein iPTM and pTM scores were recorded but not used as hard filters; all AF3 complexes, irrespective of global confidence scores, were carried forward into the NES-focused structural analysis.

### Model analysis

To generate a ranked atlas of NESs across AF3 models, we developed an unsupervised scoring matrix grounded in the structural representation of the NES helix. For each XPO1-cargo complex, we defined the hydrophobic pockets P0-P4 of the H11-H12 groove by mapping 6CIT-derived pocket centroids onto the AF3 XPO1 via structural alignment, thereby enforcing a consistent pocket frame across all cargos without imposing sequence motifs on the D-chain. Within the cargo, we then identified short α-helical segments in the vicinity of the groove and, for each helix, used side-chain centroids of hydrophobic residues (L, I, V, F, M) to assign at most one anchor per pocket within a fixed distance cutoff to the corresponding centroid. This yielded, for every model, a pocket map specifying which cargo residues occupied P0-P4, together with local geometry (anchor-pocket distances, helix tilt) and contacts to XPO1 residues in the gate-keeper region around E540 and K568/K579 (including neighboring lysines K525 and K533).

These measurements were combined into two numerical scores. The Base_Score (0-16) summarizes structural engagement of the helix with the groove: each occupied pocket contributes 1 point, with a bonus for complete P0-P4 occupancy, and additional contributions reflect anchor-pocket geometry, contact density to H11-H12, compatibility with the gate-keeper region, and the mean pLDDT of the NES core and its flanks. The Reproducibility_Score (0-5) captures model-to-model consistency by clustering candidate segments across the five AF3 models by residue overlap and pocket usage and rewarding solutions that recur with similar pocket maps and orientations. Their sum, the Model_Rank score (0-21), was used to prioritize NES segments per cargo; for each protein we report per-model scores and a consensus call (top-scoring NES, pocket map, spacing pattern, orientation, and gate-keeper contact profile) that forms the basis for downstream classification and analysis.

### NES groove plots and 2D NES/domain maps

To visualize NES geometry and context, AF3 models were converted into two complementary 2D representations. Groove plots summarize pocket occupancy and orientation at residue resolution. For each top-scoring NES, we constructed a pocket-position matrix in which residues in the NES window are arrayed along one axis and pockets P0-P4 along the other; entries reflect how often a residue is seated in each pocket across the five AF3 models. Canonical anchors (L, I, V, F, M) are highlighted, and non-canonical pocket occupants are marked separately. Groove plots were generated in Python using NumPy/pandas and Matplotlib.

In parallel, 2D NES/domain maps locate each NES along the primary sequence and embed it within the protein’s architectural context. For each UniProt entry, we assembled a linear track containing predicted intrinsic disorder (MetaPredict), domain boundaries from SMART and TED, and per-residue pLDDT from AlphaFold. NES windows corresponding to high Model_Rank scores were overlaid as colored boxes on this axis. These maps were produced in Python/Matplotlib for batch generation and, in some cases, imported into MATLAB for final layout. Together, groove plots and NES/domain maps convert 3D AF3 complexes into interpreTable S2D readouts that capture pocket usage, spacing, and domain context.

### Iterative NES classification

All scored NES candidates were classified using an iterative, unsupervised framework that integrates sequence motifs, structural orientation, and pocket occupancy. First, a sequence-based classifier used regular expressions to encode canonical NES families (1a-1d, 2, 3, 4) and their reverse counterparts with explicit inter-anchor gaps and simple chemistry gates (Φ or A/T at P1/P2 with restrictions; strict Φ at P3/P4; proline in the Φ3XXXΦ4 gap for Class-4). Palindromic forward/reverse matches were handled by a “palindromic policy” that designates a primary call and flags ambiguity, providing a structure-independent baseline for canonical matches.

Next, we incorporated orientation and pocket usage from the groove analysis. Anchors seated in P0-P4 were ordered in a forward frame and used to compute oriented inter-anchor gaps, which were mapped onto canonical families (five-anchor Classes 1a-1d and 2; Class-3 and its truncated variants; Class-4 with its characteristic P3-P4 geometry), with reverse solutions labeled “-R”. Chemistry gates were reinforced at P1-P4, and per-model class calls were aggregated into a per-cargo consensus; unresolved or conflicting cases were labeled Ambiguous. Finally, we extended this orientation-aware framework to include internal-skip and “like” solutions. Motifs that bypassed one pocket and re-engaged the groove downstream, while preserving a coherent pocket register and contact network, were assigned to Class-5, and spacing patterns that matched canonical families but relaxed one or more chemical or spacer rules were labeled as “_L” subclasses (e.g., 1a_L, 2_L) with an explicit Like_Reason. Sequence-based classifications were retained as an orthogonal check against the structure-aware assignments.

### NES validation

Selected NES candidates were validated using a live-cell reporter in which the candidate NES was fused to an SV40 NLS-GFP (or NLS-mCherry) cassette under a CMV promoter. Reporter constructs were synthesized and cloned into mammalian expression vectors (Twist Bioscience). HEK293T cells (authenticated, mycoplasma-negative) were maintained in DMEM supplemented with 10% FBS at 37 °C and 5% CO₂, plated on #1.5 glass-bottom dishes, and transfected 18-24 h before imaging. XPO1 dependence was tested by treatment with leptomycin B (10-20 nM, ∼2 h). Cells were fixed, stained with DAPI, and imaged by confocal microscopy, and NES activity was scored based on nuclear versus cytoplasmic distribution of the fluorescent reporter in the presence and absence of leptomycin B.

### NLS-NES juxtaposition analysis

To identify NLS motifs arranged in juxtaposition to AF3-identified NESs, we searched a ±50-amino-acid window around each NES along the primary sequence. Within this window, we scanned for classical basic NLS patterns: monopartite NLSs were defined as short lysine/arginine-rich stretches (K(2-5) or R(2-5)), and bipartite NLSs as two basic clusters separated by a flexible spacer, (KR)(2-5)-X(2-10)-(KR)(2-5). Overlapping hits were merged, and for each NES-containing protein we recorded the presence, type, and position of each NLS relative to the NES (N- or C-terminal within the ±50-residue window), which was then used to quantify juxtaposed NES-NLS architectures across the exportome.

### Limitations of the study

Several limitations of this work should be noted. First, although our structure-aware classifier recovers and refines canonical NES families and adds new Class-5 and “_L” subclasses, a substantial fraction of high-scoring NES candidates remains unassigned to any defined class, including the modeled NES from experimentally validated Cyclin B1. Many of these non-canonical solutions show reproducible pocket usage and clear helix-groove engagement, but fall outside our current spacing and chemistry rules. In this version of the atlas, we therefore leave these motifs “undefined” rather than over-interpreting them, and further experimental work—both functional NES assays and high-resolution structural studies—will be required to determine which of these represent bona fide new NES classes versus structural or modeling artifacts.

Second, our analyses rely entirely on AF3 models, which, despite their high overall accuracy, remain predictions. AF3 operates in a single-structure, low-crowding regime and does not fully capture conformational dynamics, partner-induced masking, phase separation, or context-dependent accessibility of NESs in vivo. Although we incorporated metal cofactors where needed (e.g., Zn-finger proteins, E3 ligases, and RQC components), these models may still not fully represent the native conformational space of the full-length protein. Thus, while NES engagement in the groove is likely to be captured with relatively high fidelity, the overall quaternary arrangement of the complex may not reflect the proper cellular context. Likewise, our focus on α-helical, hydrophobic pocket engagement, centered on the H11-H12 repeats of XPO1, means that atypical export interfaces—if they exist—are underrepresented or missed and will be evaluated in the future.

## Data Availability

Exportome Atlas, a searchable companion web portal for this study, is available at nuclear.exportome.org and enables browsing, visualization, and download of structure-resolved NES annotations, juxtaposed NLS features, groove plots, and NES location plots associated with the current preprint release. The portal and associated dataset may be updated in future versions and will remain publicly available upon publication.

## Acknowledgement

This work was supported by an R01 award from the National Institute of Arthritis and Musculoskeletal and Skin Diseases (NIAMS) to C.K. (R01AR073906) and by an Institutional Development Award (IDeA) from the National Institute of General Medical Sciences of the National Institutes of Health under grant number P20GM103436 (D.B.). The authors would also like to thank Mrs. Elizabeth Pelphrey, coordinator of the Math, Science, Technology, and Engineering (MSTC) program at Paul Lawrance Dunbar High School, in the Fayette County Public Schools (FCPS) system, for facilitating the community science aspect of this project.

