## Supplemental Figures for "A Deep-Learning Atlas of XPO1-Mediated Nuclear Export at Proteome Scale"

A. XPO1 (1071 aa) — HEAT repeats (A=blue, B=green)

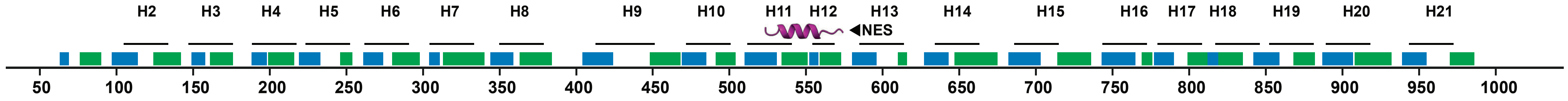


B.

| Class | Sequence Pattern |
|---|---|
| 1a | Φ1XXXΦ2XXΦ3XΦ4 |
| 1b | Φ1XXΦ2XXΦ3XΦ4 |
| 1c | Φ1XXXΦ2XXXΦ3XΦ4 |
| 1d | Φ1XXΦ2XXXΦ3XΦ4 |
| 2 | Φ1XΦ2XXΦ3XΦ4 |
| 3 | Φ1XXΦ2XXXΦ3XXΦ4 |
| 4 | Φ1XXXΦ2XXΦ3XXXΦ4 |
| 1a-R | Φ1XΦ2XXΦ3XXXΦ4 |
| 1b-R | Φ1XΦ2XXΦ3XΦ4 |
| 1c-R | Φ1XΦ2XXXΦ3XXXΦ4 |
| 1d-R | Φ1XΦ2XXXΦ3XΦ4 |

C. PDB: 3GB8 Snuportin-NES

AF3.0:Model_4 Snuportin-NES

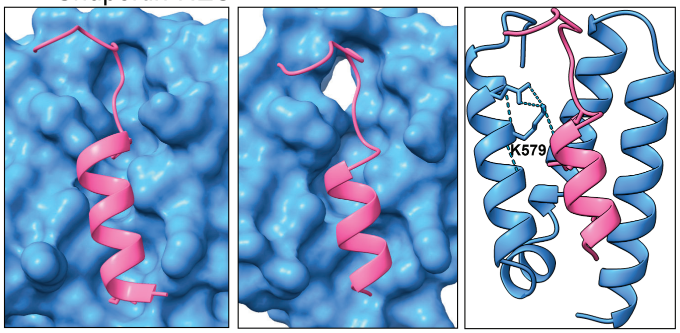


Snuportin-NES: [1]MEELSQALASSFSVS[16]

Class-1c

D. PDB: 5HDF hRioK2-NES

AF3.0:Model_4 hRioK2-NES

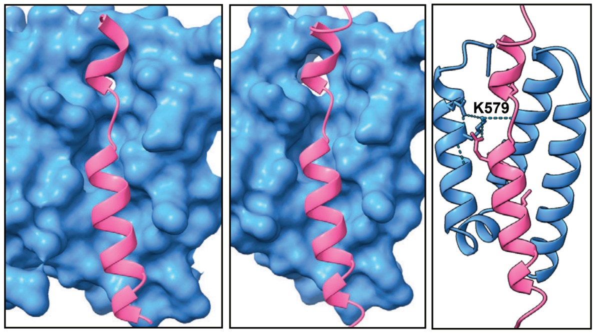


hRioK2-NES:[384]ESADARSFEMTEFNQALEEIKGQV[407]

Class-1a-R

E.

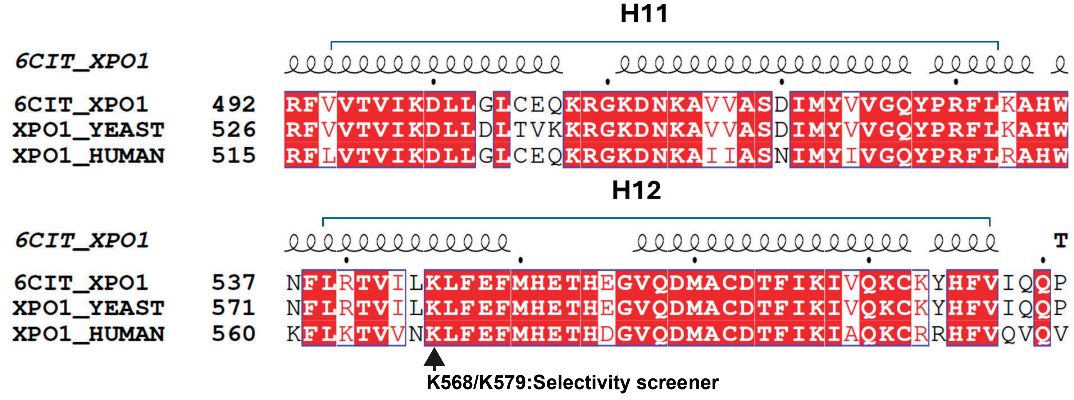


Figure S1

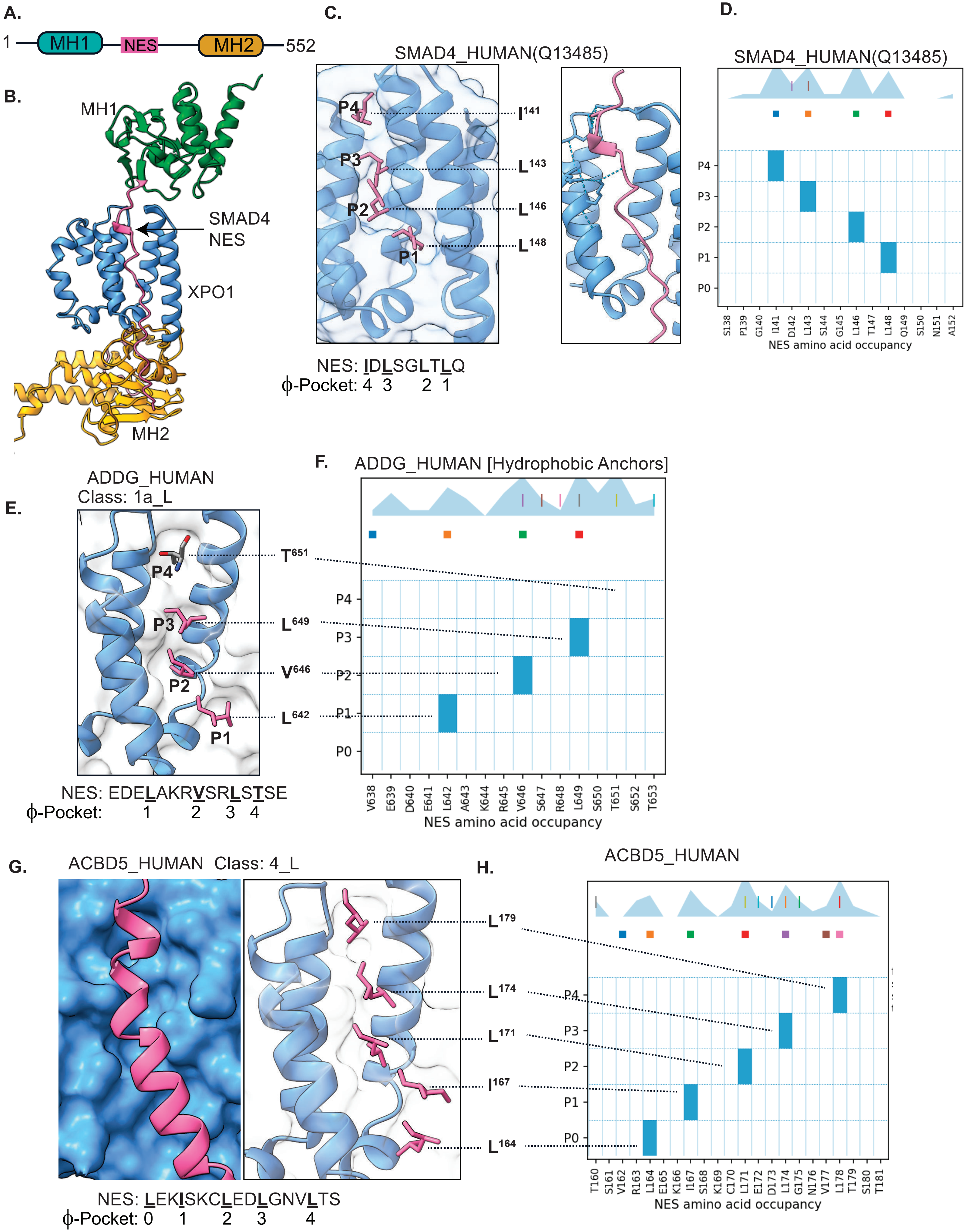
A.
1
MH1
NES
MH2
552
B.
MH1
SMAD4
NES
XPO1
MH2
C.
SMAD4_HUMAN(Q13485)
P4
P3
P2
P1
I141
L143
L146
L148
NES: IDLSGLTLQ
ϕ-Pocket: 4 3 2 1
D.
SMAD4_HUMAN(Q13485)
P4
P3
P2
P1
P0
S138 P139 G140 I141 D142 L143 S144 G145 L146 T147 L148 Q149 S150 N151 A152
NES amino acid occupancy
E.
ADDG_HUMAN
Class: 1a_L
P4
P3
P2
P1
T651
L649
V646
L642
NES: EDELAKRVSRLSTSE
ϕ-Pocket: 1 2 3 4
F.
ADDG_HUMAN [Hydrophobic Anchors]
P4
P3
P2
P1
P0
V638 E639 D640 E641 L642 A643 K644 R645 V646 S647 R648 L649 S650 T651 S652 T653
NES amino acid occupancy
G.
ACBD5_HUMAN Class: 4_L
L179
L174
L171
I167
L164
NES: LEKISKCLEDLGNVLTS
ϕ-Pocket: 0 1 2 3 4
H.
ACBD5_HUMAN
P4
P3
P2
P1
P0
T160 S161 V162 R163 L164 E165 K166 I167 S168 K169 C170 L171 E172 D173 L174 G175 N176 V177 L178 T179 S180 T181
NES amino acid occupancy

Figure S3

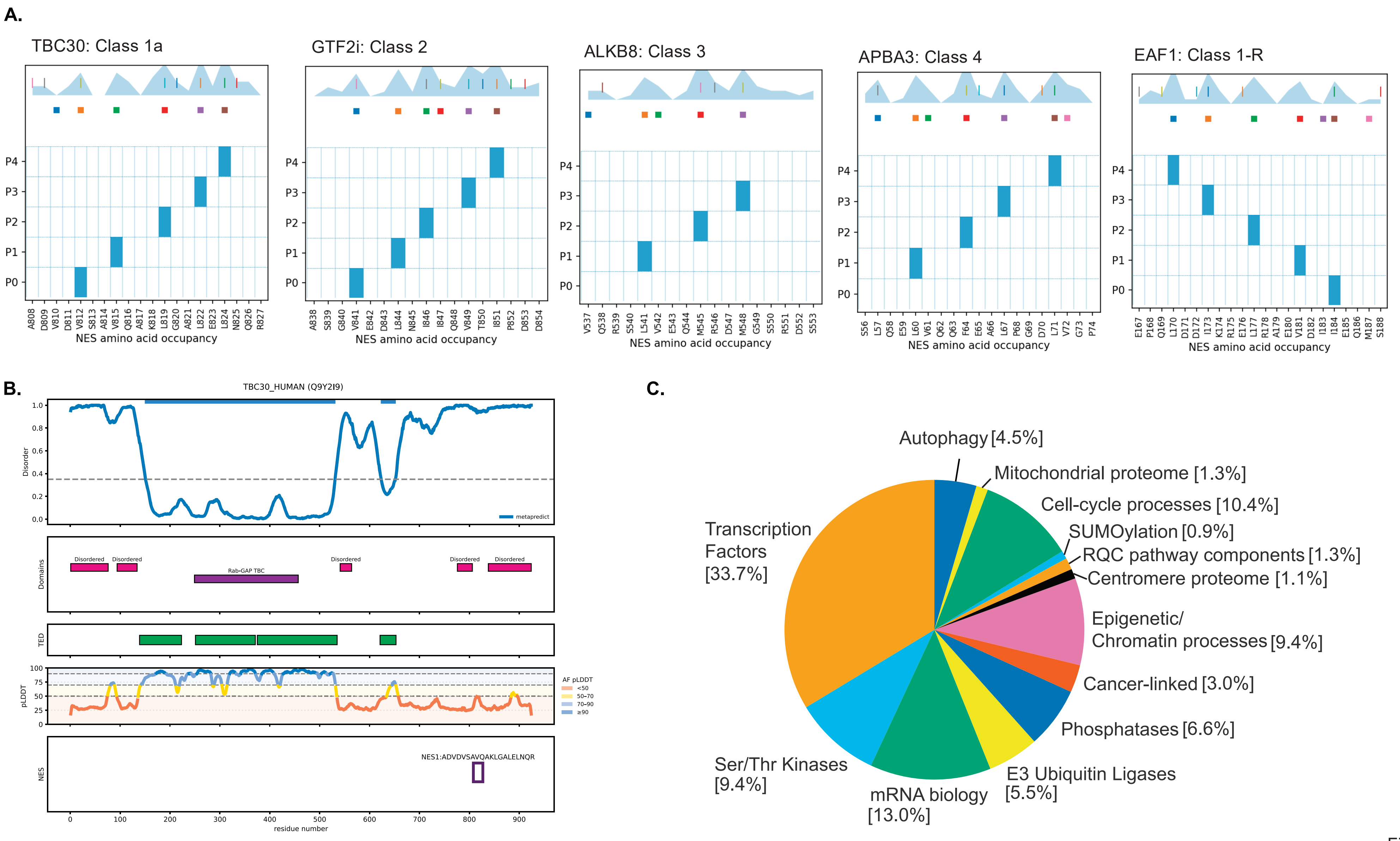
A.
TBC30: Class 1a
GTF2i: Class 2
ALKB8: Class 3
APBA3: Class 4
EAF1: Class 1-R
P4
P3
P2
P1
P0
NES amino acid occupancy
A808 D809 V810 D811 V812 S813 A814 V815 Q816 A817 K818 L819 G820 A821 L822 E823 L824 N825 Q826 R827
A838 S839 G840 V841 E842 D843 L844 N845 I846 I847 Q848 V849 T850 I851 P852 D853 D854
V537 Q538 R539 S540 L541 V542 E543 Q544 M545 R546 D547 M548 G549 S550 R551 D552 S553
S56 L57 Q58 E59 L60 V61 Q62 Q63 F64 E65 A66 L67 P68 G69 D70 L71 V72 G73 P74
E167 P168 Q169 L170 D171 D172 I173 K174 R175 E176 L177 R178 A179 E180 V181 D182 I183 I184 E185 Q186 M187 S188
B.
TBC30_HUMAN (Q9Y2I9)
Disorder
metapredict
Domains
Disordered
Rab-GAP TBC
TED
pLDDT
AF pLDDT
<50
50–70
70–90
≥90
NES
NES1:ADVDVSAVQAKLGALELNQR
residue number
C.
Transcription Factors [33.7%]
Autophagy [4.5%]
Mitochondrial proteome [1.3%]
Cell-cycle processes [10.4%]
SUMOylation [0.9%]
RQC pathway components [1.3%]
Centromere proteome [1.1%]
Epigenetic/ Chromatin processes [9.4%]
Cancer-linked [3.0%]
Phosphatases [6.6%]
E3 Ubiquitin Ligases [5.5%]
mRNA biology [13.0%]
Ser/Thr Kinases [9.4%]

Figure S2
